# Surpassing Light Inhomogeneities in Structured-Illumination Microscopy with FlexSIM

**DOI:** 10.1101/2023.12.20.572677

**Authors:** Emmanuel Soubies, Alejandro Nogueron, Florence Pelletier, Thomas Mangeat, Christophe Leterrier, Michael Unser, Daniel Sage

**Affiliations:** IRIT, Université de Toulouse, CNRS, Toulouse, France; Centre de Biologie Integrative, Toulouse, France; Biomedical Imaging Group, EPFL, Lausanne, Switzerland; Aix Marseille Université, CNRS, INP, NeuroCyto, Marseille, France; LITC Core Facility, Université de Toulouse, CNRS, Toulouse, France

**Keywords:** Super-resolution, Structured-Illumination Microscopy, Image Reconstruction, Computational Imaging, Artifact Reduction, Patterns Estimation

## Abstract

Super-resolution structured-illumination microscopy (SIM) is a powerful technique that allows one to surpass the diffraction limit by up to a factor two. Yet, its practical use is hampered by its sensitivity to imaging conditions which makes it prone to reconstruction artifacts. In this work, we present FlexSIM, a *flexible* SIM reconstruction method capable to handle highly challenging data. Specifically, we demonstrate the ability of FlexSIM to deal with the distortion of patterns, the high level of noise encountered in live imaging, as well as out-of-focus fluorescence. Moreover, we show that FlexSIM achieves state-of-the-art performance over a variety of open SIM datasets.

## 1. Introduction

Since the seminal works of Heintzmann, Cremer [1] and Gustafsson [2], super-resolution structured-illumination microscopy (SIM) has become increasingly popular [3– 7]. Among super-resolution fluorescence microscopy techniques [8–10], it is one of those that offer the best tradeoff between spatial and temporal resolution [11–13]. Moreover, it does not require specific sample preparations, offers high photon efficiency, and supports multicolor imaging [3, 14, 15].

SIM is a prime example of *computational microscopy* that combines optics and numerical reconstruction so as to surpass the diffraction limit. Specifically, a set of structured illuminations is exploited to shift high-frequency components of the imaged sample within the bandpass of the optical system. Then, through dedicated algorithms, this highfrequency information is extracted from acquired data and used to generate an image with extended resolution. From a numerical standpoint, the importance and popularity of SIM can be measured by the growing number of reconstruction software packages [16]. These include, among open-source packages: SIMToolbox [17]; FairSIM [18]; OpenSIM [19]; Hessian-SIM [20]; DL-SIM [21]; HiFi-SIM [22]; ML-SIM [23]; JSFR-SIM [24]; Direct-SIM [25]; rDL-SIM [26]; or Open-3DSIM [27].

In conventional SIM, sinusoidal illumination patterns are used, which allows one to improve the resolution of widefield microscopy by up to a factor two [1, 2]. Yet, several extensions of this conventional SIM setup have been proposed. They undertake to improve the resolution even further, to image thick samples, to reduce acquisition time, to simplify acquisition protocols, or to reduce background fluorescence [4–6]. For instance, a resolution improvement beyond a factor two is theoretically achievable if the fluorescence emission can be made to depend nonlinearly on the illumination [28–31]. Moreover, it has been shown that the use of random (speckle) illuminations improves the imaging of thick samples [32–35]. In another vein, SIM has been combined with total internal reflection fluorescence (TIRF) [36– 38] or grazing incidence [39] illuminations. These limit the excitation to a few hundred nanometers above the coverslip, which strongly reduces out-of-focus fluorescence.

Unfortunately, SIM is particularly prone to reconstruction artifacts, which hinders its practical use. This has led some researchers to characterize and classify typical SIM artifacts [40–42]. Other researchers have dedicated their work to the development of i) detailed protocols for sample preparation [43] and system calibration [40, 44]; ii) numerical tools to assess image quality [45]; and iii) guidelines to best take advantage of reconstruction software packages [38, 40–42].

The quest for artifact-free SIM reconstruction is currently a very active area of research, as evidenced by the recent surge in numerical strategies such as the shaping of the reconstruction point-spread function (PSF) or optical transfer function (OTF) into an ideal form to reduce commonly seen artifacts [22, 46], the deployment of rolling reconstructions to mitigate motion artifacts in life imaging [20], the estimation and filtering of background fluorescence [47–49], the use of blind-reconstruction approaches [50–52], as well as the control of noise to remove structured-noise artifacts [53]. Finally, it is worth mentioning that deep neural networks have also been designed and trained to reduce artifacts in SIM reconstructions [54, 55].

This situation (along with the difficulties we experienced ourselves upon striving to significantly reduce reconstruction artifacts for challenging TIRF-SIM data using existing methods), motivated the development of FlexSIM. The promise of FlexSIM, for *flexible* SIM reconstruction, is to provide reliable SIM reconstructions for a variety of SIM data, going from “ideal” ones acquired under standardized protocols and configurations, to ones obtained under more challenging settings and more prone to reconstruction artifacts.

## 2. Materials and Methods

### A. Foundations of FlexSIM

FlexSIM builds upon three pillars, highlighted in Figure 1.

**Fig. 1.**
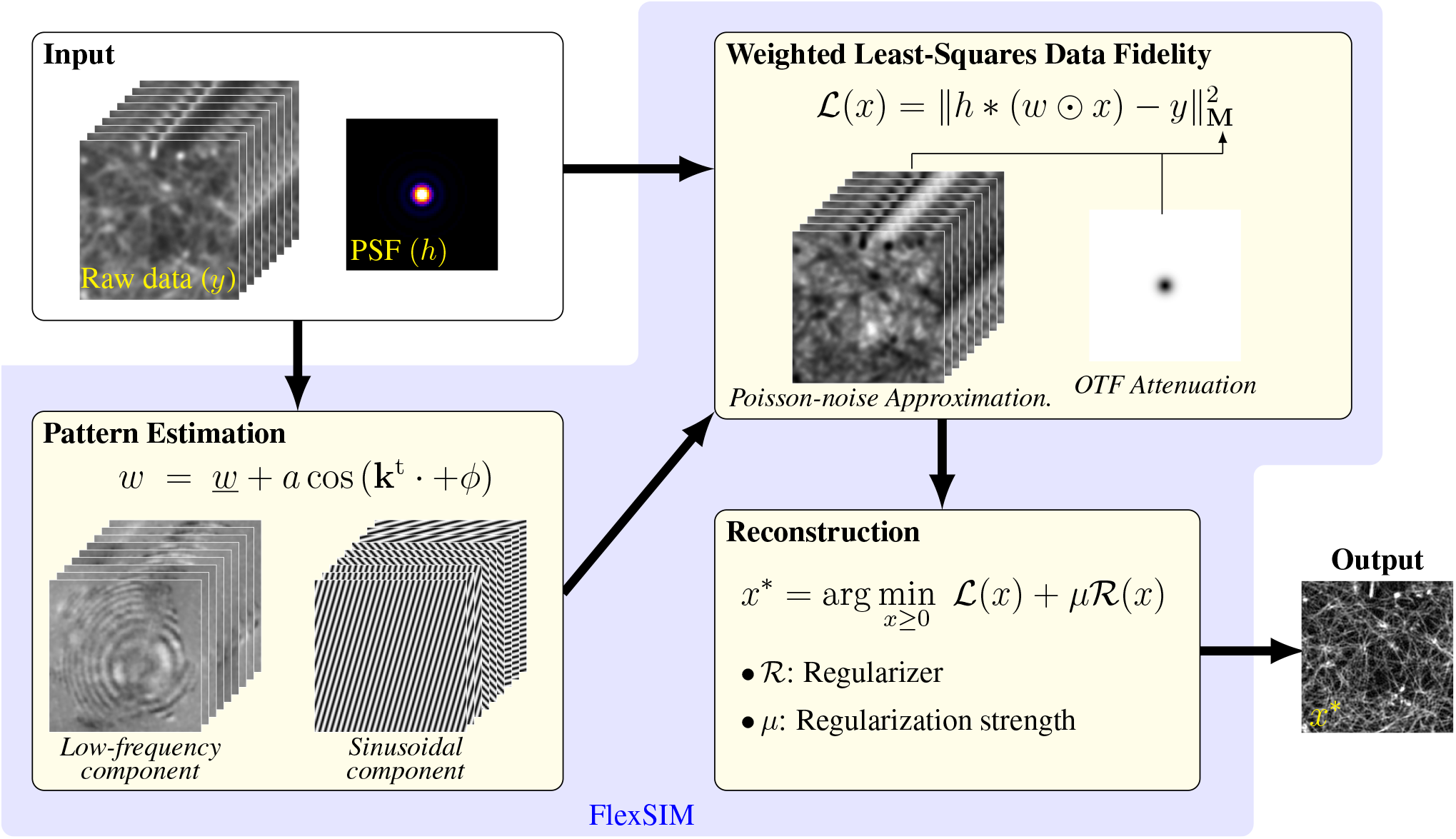
Flowchart of FlexSIM. In addition to the ideal sinusoidal illuminations, low frequency pattern components 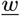 are estimated from raw data. For reconstruction, a weighted least-squares data-fidelity term is considered. The weighting operators **M** are built so as to account for both out-of-focus signal (similarly to OTF attenuation) and shot-noise. Finally, reconstruction can be performed with a variety of regularizers ℛ and optimization algorithms.

#### 1) Improved modelling

We propose the use of a more realistic model to represent the experimental illumination patterns. Specifically, we consider

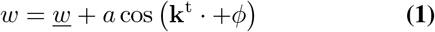

where <u>*w*</u> represents a low-frequency component (to be estimated from data) that refines the simple model of a purely sinusoidal SIM pattern with amplitude *a*, wave vector **k**, and phase *ϕ*. This refinement turned out to be essential to treat challenging TIRF-SIM data, as shown in Figure 2. Then, reconstruction is performed using a weighted-leastsquares data-fidelity term that allows us to account for both out-of-focus signal (similarly to OTF attenuation in standard Wiener-based reconstruction) and shot noise.

**Fig. 2.**
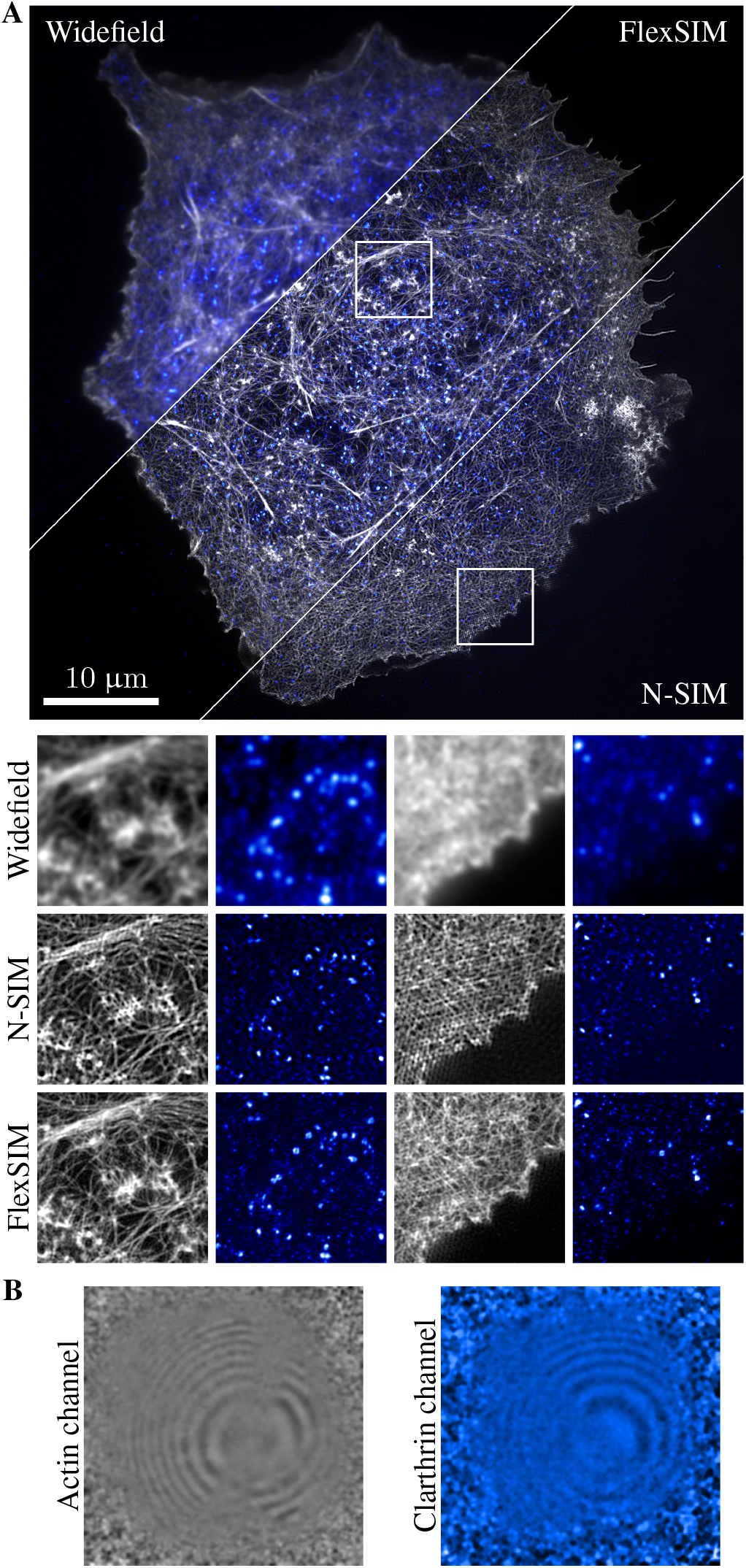
FlexSIM reconstruction of challenging TIRF-SIM data. Actin network (gray) and clathrin (blue) in COS-7 cells imaged using the Nikon N-SIM-S system in TIRF-SIM mode. Raw images are of size (1024 *×* 1024) and reconstructions are (2048 *×* 2048). **A)** Comparisons of reconstructions obtained with FlexSIM and the N-SIM software. **B)** Low-frequency patterns components estimated by FlexSIM. Reconstructions of similar datasets are presented in Figures S7 and S8.

#### 2) Auto-calibration of patterns

In FlexSIM, pattern phases and orientations are estimated through the optimization of the new criterion in (4), which we tackle in a two-step procedure. The criterion is first evaluated on a grid of candidate parameters over which optimal ones are selected. This grid-based initialization, which can be efficiently performed through cross-correlation computations, somehow recovers the widespread approach of the seminal work [2], without the need to unmix frequency components. The proposed approach can thus work with uneven phase shifts. This initial estimation is then refined (off-the-grid) through a gradient descent over the proposed criterion. Regarding the estimation of the low-frequency component <u>*w*</u>, we proposed in FlexSIM an efficient method to estimate it (one per SIM image) directly from the data (see Section B.4).

#### 3) Advanced reconstruction

Lastly, FlexSIM benefits from the GlobalBioIm framework [56] which offers a variety of regularizers and optimization algorithms. By default, first-order Tikhonov, smoothed total-variation and good-roughness regularizers are proposed in FlexSIM. Optimization is performed with a quasi-Newton approach.

We provide in Sections B and C the methodological details behind each step of the FlexSIM pipeline.

**Notations**.. We write vectors as bold lowercase letters (e.g., **x, k**) and their transpose as **x**^t^, **k**^t^. The *m*th component of a vector **x** ∈ ℝ^*M*^ is *x*_*m*_. We denote the Fourier transform of a function *v* (lowercase) by it uppercase counterpart *V*. The symbols * and *0* stand for convolution and pointwise multiplication operations, respectively. The complex conjugate of *a* ∈ ℂ is denoted ā. We denote by ⟨·, ·⟩ the standard Hermitian inner product, the conjugation being applied to the first argument by convention.

### B. Pattern Estimation Module

#### B.1. The Art of Pattern Estimation

An accurate pattern estimation is essential to limit reconstruction artifacts. The vast majority of existing approaches work under the assumption that the spatial phases shifts of patterns sharing the same orientation are regularly spaced within [0, 2*π*]. This allows for a simple separation of the Fourier components contained in the raw SIM images from the sole knowledge of relative phases. Then, the maximization of the cross-correlation between these extracted Fourier components leads one to estimate the orientations of the patterns, as well as their frequencies and phase offsets [2, 18, 57]. While this strategy proved to be very efficient to estimate pattern orientations and frequencies, it may not provide a sufficiently accurate estimation of phases. As such, several works have been dedicated to phase estimation, assuming that orientations and frequencies are known and that phases are not necessarily regularly spaced [58–61]. Finally, there exist alternative approaches that exploit either Prony’s annihilation property [62], or the low-rank nature [63] of SIM illuminations.

#### B.2. Patterns Estimation with FlexSIM

Let 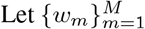 be *M* illumination patterns having the same orientation but different phases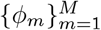. In conventional 2D-SIM, patterns are generated from the interference between two beams and are modeled as

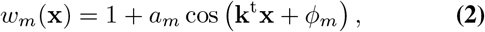

where *a*_*m*_ *>* 0 is the modulation contrast, **x** ∈ ϕ^2^ the spatial variable, and **k** ∈ ℝ^2^ the modulation light wave vector. The associated (noiseless) SIM data 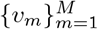 are then given by

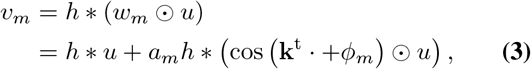

where *u* represents the sample and *h* the PSF of the optical system. In the sequel, we denote by 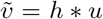 the widefield image, which we assume to be accessible. When the phases are equally spaced, we have that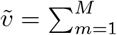 *v_m_*. Otherwise, the widefield image can be acquired together with the SIM stack.

To drop the dependence on the sample *u*, which is unknown, we make the approximation that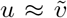. Then, pattern parameters can be estimated through the minimization with respect to **a, ϕ**, and **k** of the criterion

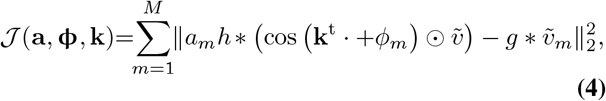

where **ϕ** ∈ [0, 2*π*)^*M*^, 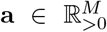, and 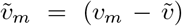 corresponds to the *m*th raw SIM data without the widefield component. In practice, to account for scaling factors, we compute 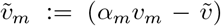 for *α*_*m*_ that minimizes the error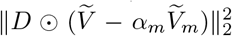, that is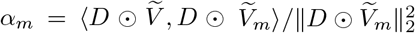. Here, *D* is a mask that allows us to compute the error between low frequencies only (e.g., a disk in Fourier). Finally, *g* is a filter introduced to mitigate the effect of the approximation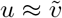. More precisely, the approximation 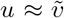 amounts to consider (in first approximation) that we can commute the convolution and multiplication operators. This generates a mismatch error between the simplified model 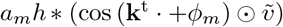 and the data 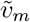 as illustrated in the third column of Figure S3. The role of *g* is thus to mitigate this mismatch (cf. last column of Figure S3). It is related to the idea of notch filtering used in other works [22].

The function 𝒥 is highly nonconvex, making its minimization a challenging task. As such, we propose a two-step procedure.

#### 1. Grid-Based Initialization

The idea here is to consider an approximation of 𝒥 that can be efficiently evalu-ated on a grid 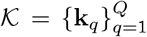 of candidate wave vec-tors, and then select **k**_init_ = min_**k**∈𝒦_ 𝒥 (**k**) ∈ 𝒦. We refer the reader to *Supplementary Note 1* for a full derivation. Similar to standard pattern-estimation approaches [2, 59], the proposed grid-based initialization relies on cross-correlation computations. Yet, it does not require us to first unmix the frequency components. The advantage is that it can work with uneven phase shifts in a non-iterative way. The main computational cost lies in the computation of *M* crosscorrelation maps. In that respect, it has links with the method proposed in [60].

#### 2. Local Off-the-Grid Refinement

The goal here is to improve the initial estimation through a local optimization of 𝒥. This allows us to obtain an estimate of the wave vector that is not constrained to be on the grid 𝒦 used at initialization. Moreover, the proposed local refinement deals with the exact function 𝒥. It proceeds by alternating between a minimization of 𝒥 with respect at first to (**a, *ϕ***), and then **k**. For the minimization over (**a, *ϕ***), we recast the problem as the resolution of *M* (2 *×* 2) systems of linear equations that can be solved efficiently. Regarding the minimization of 𝒥 over **k**, we derive a closed-form expression of the gradient and deploy gradient-descent steps. Full details are provided in *Supplementary Note 1*.

#### B.3. Numerical Validation

We report in Figure S4 a series of numerical experiments over simulated data. As expected, we observe that errors decrease when the level of noise decrease, that is when the maximal expected number of photons (MEP) increases. For all noise levels, the wave vector is estimated with a precision that is beyond a tenth of pixel and reaches a hundredth of pixel for low noise levels. For the estimation of phases, errors vary from less than 10° for high noise levels to less than 1° when the noise decreases (Figure S4-**C**). These are similar to the errors obtained with state-of-the-art methods [59–61].

An important outcome is that the same accuracy is reached with or without the assumption that the phases are equally spaced. However, challenging real data such as those acquired in a TIRF-SIM mode do benefit from the assumption that the phases are equally spaced.

Finally, let us comment on how oversampled data influence the quality of the cross-correlation map (initialization step) (Figure S4-**B**). As expected, pattern-estimation errors decrease as the oversampling increases. Yet, even with a sixteen-fold oversampling, the cross-correlation based initialization does not reach the accuracy obtained after our refinement step. This is especially true for MEP larger than 10, which corresponds to typical noise levels encountered in practice. This highlights the relevance of the proposed local refinement step. Moreover, large oversampling factors significantly increase the memory usage and computational time.

For instance, for the (512 *×* 512) image used in Figure S4, the cross-correlation computation with a sixteen-fold over-sampling is six times slower than the two-step approach: initialization (with twofold oversampling) plus local refinement (∼30 s vs ∼5 s). As such, given that exceeding a twofold oversampling for the initialization does not lead to a significant gain in the refinement step, the oversampling factor is fixed to 2 in FlexSIM.

#### B.4. Estimation of Pattern Low-Frequency Component

An important feature of FlexSIM is that it allows us to consider patterns of the form

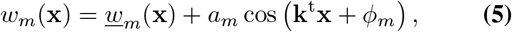

where <u>*w*</u>_*m*_ is a low-frequency component. This model proved to be crucial to obtain meaningful reconstructions for the TIRF-SIM data of Figures 2, 3, S6–S9.

**Fig. 3.**
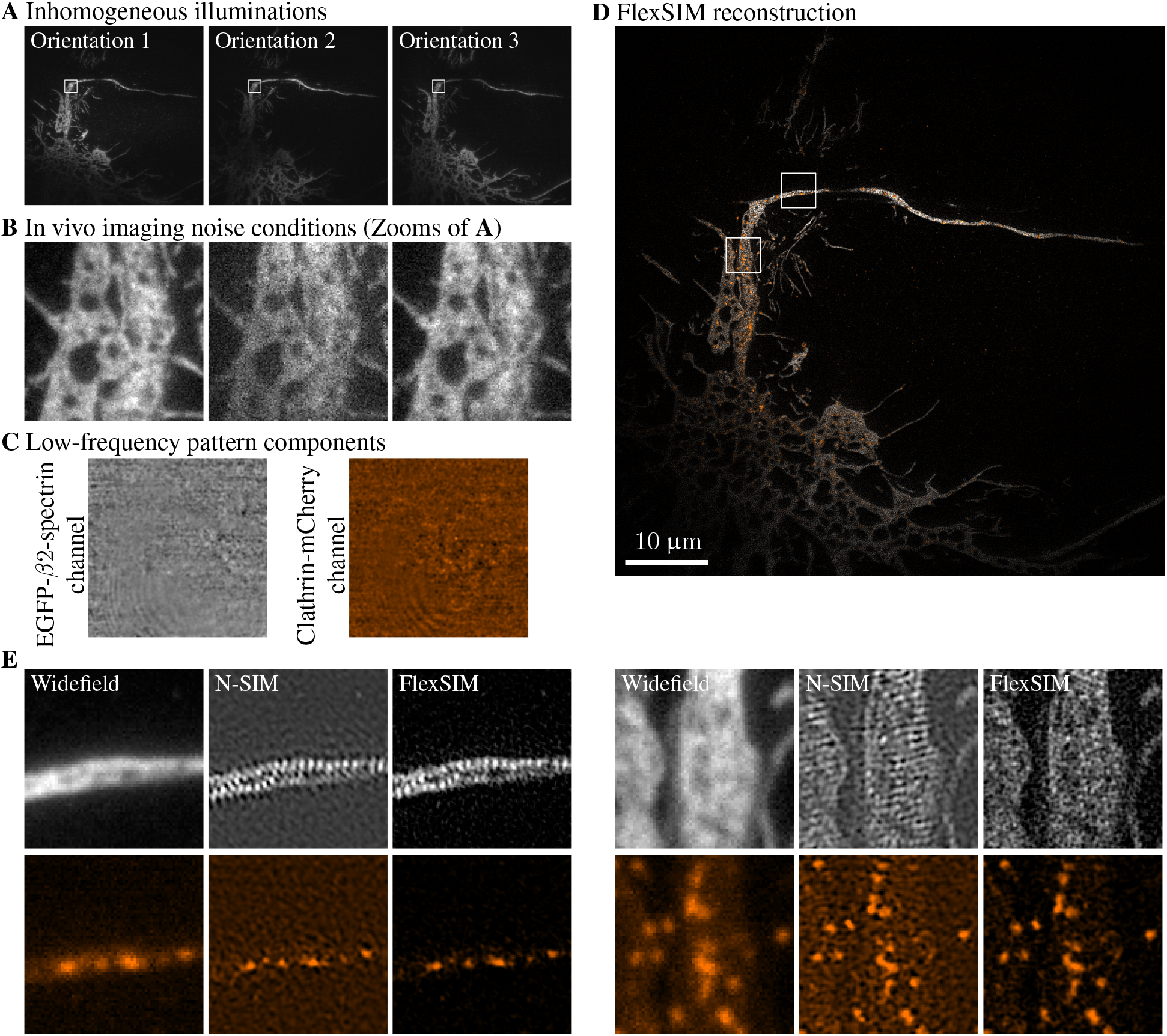
FlexSIM in live TIRF-SIM imaging conditions. Clathrin-mCherry (orange) and *β*2-spectrin expressed by a living neuron (gray). An 18-frame acquisition is performed with the Nikon N-SIM-S system in TIRF-SIM mode. Raw images are of size (1024 *×* 1024) and reconstructions are (2048 *×* 2048). **A**) Raw SIM data exhibit inhomogeneous illumination as well as **B**) high noise levels. **C**) Lowfrequency pattern component estimated by FlexSIM. **D**) FlexSIM reconstruction (the full temporal stack available in Video S1). **E**) Zooms for the comparison of FlexSIM and N-SIM reconstructions.

Regarding the estimation of <u>*w*</u>_*m*_, we deploy a simple and fast method. It relies on the fact that, given a low pass filter *f*, we get from (3) and (5) that

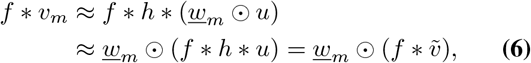

where we recall that 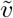 stands for the widefield image. From these approximations, we propose to use

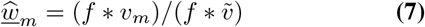

as an estimate of <u>*w*</u>_*m*_ (with pointwise division). This approach is illustrated in Figure S5.

### C. Image Reconstruction Module

#### C.1 From Direct Wiener Inversion to Iterative Reconstruction Approaches

Most SIM reconstruction methods are variants of a direct Wiener inversion [2]. For standard sinusoidal illuminations, one can easily establish and solve a closed system of equations to unmix the Fourier components of SIM data and place them back to their right location in the Fourier domain. Then, a final Wiener filtering can be used to invert the effect of the OTF and limit the amplification of noise. More evolved deconvolution approaches, for instance those that account for Poisson noise, can also be used in this final step [64]. Alternatively, the consideration of a more general variational framework allows for the use of arbitrary (non-sinusoidal) illuminations [65], a reduced number of images [66], advanced regularization terms [47, 66–68], additional out-of-focus planes in the model [47, 48], or blind reconstruction (i.e., joint pattern estimation and reconstruction) [50, 51].

#### C.2. Image Reconstruction with FlexSIM

We consider a variational reconstruction framework. Specifically, given *P* SIM raw images 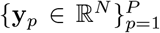 composed of *N* pixels and associated with the (discretized) patterns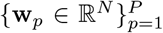, we compute the reconstructed image **û** ∈ R^4*N*^ (over a twice finer grid) as

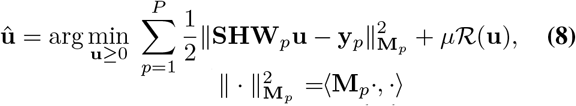

where **S** ∈ ℝ^*N×*4*N*^ is a twofold (in each dimension) downsampling operator, **H** ∈ ℝ^4*N×*4*N*^ a convolution operator defined from the system PSF, and **W**_*p*_ = **diag**(**w**_*p*_). Data fidelity is enforced using the “weighted” *L*_2_-norm 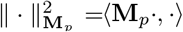, where **M**_*p*_ ∈ ℝ^*N×N*^ is defined so as to account for Poisson noise and background/out-of-focus fluorescence (details hereafter). Finally, ℛ is a regularization term and *µ >* 0 a parameter that controls the tradeoff between data fidelity and regularization.

The FlexSIM reconstruction module is implemented within the GlobalBioIm framework [56]. As such, a variety of regularizers ℛ and optimization algorithms to solve (8) are available. By default, first-order Tikhonov, smoothed total-variation [69] and good-roughness [70] regularizers are proposed in FlexSIM. They all lead to a differentiable objective function in (8) that can be minimized through the second-order variable-metric limited-memory-bounded algorithm [71] belonging to the family of L-BFGS methods. For all the experiments reported in this work, we considered the first-order Tikhonov regularizer which leads to faster computation and always provided very satisfactory results even with challenging data. We attribute the fact that such a simple regulariser is sufficient to the redundancy that exists in the 9 SIM raw images [72]. Indeed, this redundancy reduces the impact of using more sophisticated regularisers.

#### C.3. Weighted L_2_-norm

In FlexSIM, we exploit the weighted *L*_2_ data-fidelity term 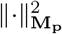 to account for both Poisson noise and out-of-focus fluorescence.

##### Poisson Noise

As in any fluorescence microscopy system, the noise in the raw SIM data is a mix of Gaussian (e.g., readout noise) and Poisson (photon-counting process of the detectors) noise. Because the associated loglikelihood is very challenging to optimize [73], standard practice is to adopt the simplifying assumption that the overall noise is a non-stationary uncorrelated Gaussian noise. This can be achieved by setting **M**_*p*_ = **diag (1***/*(**y**_*p*_ + *σ*^2^)) (component-wise division), where *σ*^2^ corresponds to the variance of the Gaussian part of the noise [67, 74].

##### Out-of-Focus Fluorescence

Out-of-focus fluorescence can be a source of severe reconstruction artifacts [40, 41]. To address this issue, reconstruction approaches that explicitly model and estimate the background signal have been proposed [47–49]. Another very common strategy is known as OTF attenuation [59]. It proceeds by attenuating the central frequencies of the OTF used in the Wiener-based final reconstruction. Basically, this is achieved through a pointwise multiplication of the OTF with a function of the form

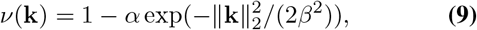

where *α >* 0 and *β >* 0 control the strength and the width of the attenuation, respectively. In this way, the frequency components that do not transmit any information about the missing cone are attenuated, thereby enhancing the components that can instead fill the missing cone (we refer the reader to the very instructive [59, Figure 7]). From a variational viewpoint such as (8), this OTF-attenuation strategy amounts to set **M**_*p*_ = **H**_Att_, a convolution operator whose kernel is given in Fourier domain by *ν* in (9). Indeed, with this choice and setting 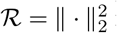 in (8), we recover the standard Wiener-based reconstruction with OTF attenuation. This particular formulation allows us to interpret OTF attenuation as a means of giving more importance to high-frequency errors than to low-frequency errors (including errors due to the background) in the data-fidelity term.

In accordance with this section, we define in FlexSIM the data-fidelity term with **M**_*p*_ a composition of **diag (1***/*(**y**_*p*_ + *σ*^2^)) and **H**_Att_.

### D. Sample Preparation

COS-7 cells were briefly extracted in Triton X100/glutaraldehyde, then fixed using glutaraldehyde, before being quenched, blocked, and stained with anti-clathrin heavy-chain primary antibodies (polyclonal rabbit ab21679, abcam), revealed with donkey anti-rabbit secondary antibodies conjugated to Alexa Fluor 647, and anti-alpha tubulin primary antibodies (monoclonal mouse clones B-5-1-2 and DM1a, Sigma), revealed with donkey anti-mouse secondary antibodies conjugated to Alexa Fluor 555. To stain actin, cells were incubated with phalloidin-Atto488 at the end of the staining procedure [75].

Regarding macrophages imaging (Figure S9), macrophages were transduced with GFP-paxillin lentiviruses (BiVic facility, Toulouse, France) for three days as previously described [34]. Macrophages were fixed with paraformaldehyde and were placed on a FluoroDish (WPI FD35-100) and immersed in PBS.

### E. TIRF-SIM Imaging and N-SIM Processing

We used a Nikon N-SIM-S microscope to image COS-7 cells stained for actin via classical two-beam TIRF-SIM. Cells were mounted in a Ludin chamber in 0.1M phosphate buffer.

The sample was illuminated using a 488 nm laser with 2 opposite beams at the periphery of the back focal plane of a 100X, 1.49 NA objective, with 9 images (3 phases X 3 orientations; 16-bit, (1024 *×* 1024) pixels at 65 nm/pixel) captured over a 50 ms exposure time by an Hamamatsu Fusion BT sCMOS camera. The raw images were then processed using the N-SIM module of the NIS Elements software, resulting in a 32-bit, (2048 *×* 2048) pixel reconstructed image at 32.5 nm/pixel.

## 3. Results

### A. FlexSIM Deals with Pattern Distortions

An important feature of FlexSIM lies in its ability to cope with illumination patterns that exhibit a nonuniform low-frequency component, as described in (1). We report in Figure 2 an example of challenging TIRF-SIM data that are affected by such a degradation. Although not clearly visible on individual raw images (Figure S5.A), slowly varying concentric rings become apparent if one switches rapidly from one image to the next. The origin of these distortions is not entirely clear. They are most likely due to reflections from the TIRF edge configuration and, to date, cannot be systematically corrected on the optical side. Fortunately, these distortions are captured perfectly by the low-frequency component <u>*w*</u> estimated by FlexSIM (Figures 2.B and S5.B). Then, by incorporating these into the reconstruction process and by reconstructing each orientation separately, FlexSIM is able to produce well-contrasted and sharp images while standard SIM-reconstructions suffer from strong grid artifacts (Figure 2.A). Even the advanced OTF shaping proposed in HiFiSIM [22] fails to avoid grid artifacts (Figure S6.B). In contrast, FlexSIM is able to significantly reduce these artefacts, albeit we can observe that some still remain in localized areas (cf. bottom left border of the cell in Figure 2.A). Finally, it is worth mentioning that the grid artifacts reappear on FlexSIM reconstructions if we ignore <u>*w*</u> or if we do not consider each pattern orientation separately (Figure S6.A). These complementary experiments strengthen the relevance and importance of the new features proposed in FlexSIM.

We report in Figures S7 and S8 two additional reconstructions over the twelve similar cell samples we acquired on the same TIRF-SIM system. One observes that, when the estimated low-frequency pattern component <u>*w*</u> is homogeneous (clathrin channel in Figure S7.C, actin and clathrin channels in Figure S8.C), no artifacts are visible in the N-SIM reconstruction as well. This reinforces the fact that the presence of a nonuniform <u>*w*</u> is at the origin of the observed artifacts. Moreover, it is worth mentioning that, even when N-SIM reconstructions do not present artifacts, the corresponding FlexSIM reconstructions exhibit better contrast and dynamics.

### B. FlexSIM Copes with Difficult Live Imaging Conditions

The TIRF-SIM data of Figure 2, which are already quite challenging, were acquired in fixed imaging conditions with high photon collection and low noise. In Figure 3, we consider even more challenging data by assessing the performance of FlexSIM in extreme live-imaging conditions. Precisely, we imaged a living neuron transfected with EGFP-*β*2-spectrin and clathrin-mCherry [76]. In addition to the presence of a low-frequency pattern component (Figure 3.C), these data present several other difficulties. In particular, they suffer from low photon collection (high noise) and strong illumination inhomogeneities that also vary according to the orientation of the pattern (Figure 3.A-B). Additionally, the imaged structure (neuron) only covers a small portion of the field of view. Lastly, marker strips spaced 190nm apart are present along the axon (horizontal structure). They are not visible on the widefield image and their spatial frequency is close to that of the grid artifacts.

The FlexSIM reconstruction of a temporal frame is presented in Figure 3.D. For comparison, the reconstruction obtained using the N-SIM software is depicted in the zooms of Figure 3.E. The full reconstructed temporal stack is provided in Video S1. While both methods are able to reveal the spectrin bands (Figure 3.E, left), they appear somehow distorted in one direction in the N-SIM reconstruction. We attribute this to the presence of grid artifacts intermingled with the actual pattern of the spectrin bands. Such grid artifacts are clearly visible in the proximal axon and dendritic region on the N-SIM reconstruction (Figure 3.E, right). Finally, FlexSIM has the remarkable ability to produce reconstructions free from residual background.

We also evaluated the performance of FlexSIM on data collected using a home-built TIRF-SIM system in Figure S9 (live macrophage expressing GFP-paxillin [35]). To illustrate the difficulty of reconstructing these data, we were unable to estimate the frequencies and phases of patterns using FairSIM [18]. While Hifi-SIM [22] performed better for this task, the best reconstruction we managed to obtain with it (or JSFR-SIM [24]) suffer from important artifacts. In contrast, FlexSIM was able to properly estimate patterns parameters and to provide a clear reconstruction.

### C. FlexSIM Achieves State-of-the-Art Performance on a Variety of Open SIM Datasets

To further demonstrate the capabilities of FlexSIM, we conducted in-depth comparisons with existing SIM reconstruction approaches. We considered twenty open 2D-SIM datasets sourced from seven publications and acquired with a diversity of SIM systems and configurations (see Table S2). Then, we benchmarked FlexSIM against methods developed in publications associated with each of these datasets. This represents seven different SIM reconstruction approaches ranging from standard Wiener-based reconstructions (and more advanced variants) all the way to deep-learning techniques. Accordingly, we could tune the parameters of each method as specified by the authors themselves, which ensures a fair comparison.

A representative subset of the comparisons we made with the most recently published reconstruction approaches is pre-sented in Figure 4. Complementary comparisons are provided in Figures S10–S13. One can see that FlexSIM performs as well as the best-in-class HiFi-SIM [22] and JSFRSIM [24] methods while outperforming other algorithms. In particular, due to the proposed weighted-least-squares data-fidelity term, including an OTF attenuation strategy, FlexSIM is able to attenuate out-of-focus signal and typical associated artifacts similarly to what is achieved by HiFi-SIM and JSFRSIM. As with the TIRF-SIM data presented in the previous paragraphs, we can appreciate the ability of FlexSIM to systematically provide well-contrasted and crisp reconstructions across the large variety of data gathered in Table S2. In addition to these extensive experiments on open real data, we report in *Supplementary Note 2* quantitative comparisons on simulated data. These results support the fact that FlexSIM not only handles very difficult data (as demonstrated in previous paragraphs), but also achieves peak performance on more standardized datasets.

**Fig. 4.**
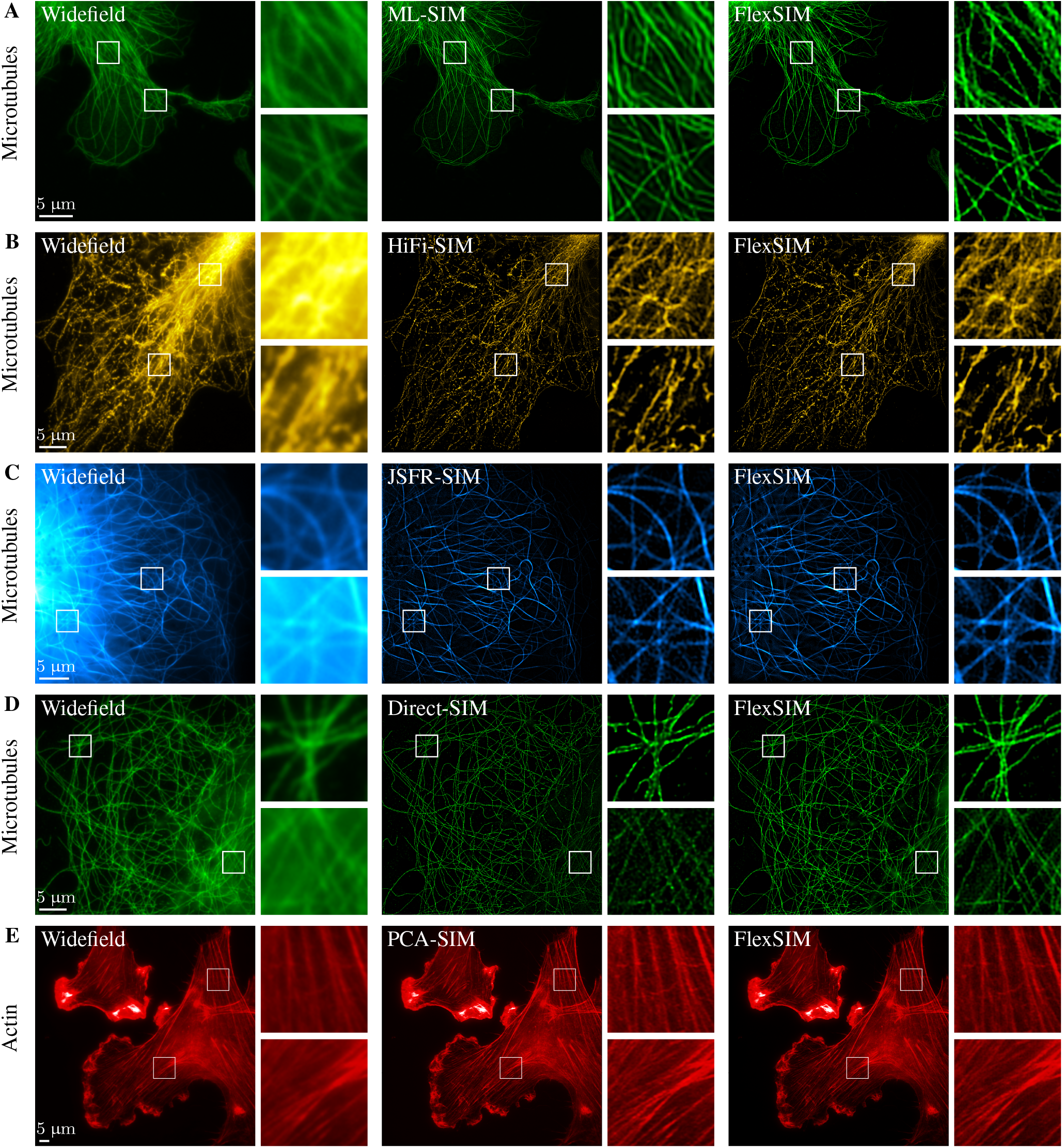
Benchmarking FlexSIM against existing software on their own datasets. A subset of the 20 open 2D-SIM datasets reported in Table S2. From left to right, columns correspond to the widefield image, the comparative reconstruction and the FlexSIM reconstruction. Other comparisons are provided in Figures S10–S13. Note that PCA-SIM [63] in **E** is a method that uses principal component analysis for the estimation of patterns parameters. The reconstruction is then performed using HiFi-SIM [22].

Finally, FlexSIM Matlab scripts for each dataset of Table S2 are publically available within the FlexSIM GitHub repository. These scripts automatically download the raw SIM data and set all FlexSIM parameters used to reproduce the results reported in the present paper for the sake of reproducibility. Beyond allowing us to assess FlexSIM performance, another motivation was to catalogue open 2D-SIM datasets and facilitate their access. We intend to update and enrich this collection of datasets as new ones are released.

## 4. Discussion

We have demonstrated experimentally the flexibility of FlexSIM to cope with a broad range of SIM data. We have included very challenging data for which no published method was known to attenuate the reconstruction artifacts as well as FlexSIM. To achieve this, we equipped FlexSIM with advanced features that include the modeling of a class of pattern distortions, the consideration of shot noise, and the attenuation of out-of-focus fluorescence. The price to pay, however, is the need for an iterative reconstruction scheme which is slower than direct Wiener-based inversion. Typically, our reconstruction of the (2048 *×* 2048) TIRF-SIM images (e.g., Figures 2 and 3) is obtained in about six minutes on a Dell Latitude computer (Intel Core i7-8650U CPU 1.90GHz × 8) with parallel computing. The reconstruction time for smaller (1024 *×* 1024) images ranges between 1 to 3 min, depending on the number of iterations. We expect that the computational time of FlexSIM can be markedly decreased if one deploys GPU computation [77].

We acquired new insights while developing FlexSIM. In particular, the estimation of pattern frequencies and phases through the formalization of this problem as the minimization of a suitable criteria helped us to clarify certain assumptions that had remained implicit to this day. This has allowed us to integrate them properly within the problem (see Figure S3). Moreover, we have highlighted the benefit of our local off-the-grid refinement over an oversampling of the data to achieve subpixel wave vector localization (see Figure S4).

Finally, FlexSIM is available as a documented opensource code with a pool of example scripts that allow one to reproduce the reconstructions reported in this work for each dataset of Table S2. Users can also deploy FlexSIM easily on any other dataset by filling a single file collecting every required parameter.

## Declarations

## Availability of data and materials

The Matlab FlexSIM code is available from the GitHub repository for the project https://github.com/esoubies/FlexSIM In addition to the core functions of FlexSIM, this repository also contains one script per dataset of Table S2 to reproduce the results of this paper.

The scripts provided in the FlexSIM GitHub repository allow for an automatic download of the datasets listed in Table S2. The TIRF-SIM data of Figures 2, 3, S7, S8, and S9 can be made available upon reasonable request.

## Authors’ contributions

D.S. and E.S. developed the core methodology of FlexSIM with M.U. inputs. A.N. and E.S. implemented FlexSIM and made all comparisons experiments with state-of-the-art SIM reconstruction approaches. F.P. prepared COS-7 cells of Figures 2, S7 and S8. C.L. imaged these samples with the Nikon N-SIM-S system in TIRFSIM mode. T.M. acquired the TIRF-SIM data of Figure S9 on a home-build system. E.S. made FlexSIM reconstructions of all these TIRF-SIM data. C.L., T.M., D.S., and E.S. analysed results. E.S. wrote the manuscript with inputs from all authors.

## Acknowledgements

The authors thanks Renaud Poincloux from Institut de Pharmacologie et de Biologie Structurale in Toulouse for the preparation of the macrophage podosomes used in Figure S9. C.L. acknowledges the INP NCIS imaging facility and Nikon Center of Excellence for Neuro-NanoImaging for service and expertise.

