## Supplementary Information for "Surpassing Light Inhomogeneities in Structured-Illumination Microscopy with FlexSIM"

**Emmanuel Soubies, Alejandro Nogueron, Florence Pelletier, Thomas Mangeat, Christophe Leterrier, Michael Unser, and Daniel Sage**

### Supplementary Note 1. Details on FlexSIM Pattern Estimation

In this note, we describe the two steps—namely *Grid-based initialization* (Section A) and *Local off-the-grid refinement* (Section B)—used to minimize $\mathcal{J}$ in (4) of the main article file, which we recall for completeness

$$\mathcal{J}(\mathbf{a}, \boldsymbol{\phi}, \mathbf{k}) = \sum_{m=1}^{M} \|a_m h * \left(\cos\left(\mathbf{k}^{\mathrm{t}} \cdot + \phi_m\right) \odot \tilde{v}\right) - g * \tilde{v}_m\|_2^2. \quad \textbf{(10)}$$

Here, $\boldsymbol{\phi} \in [0, 2\pi)^M$, $\mathbf{a} \in \mathbb{R}^M_{\geq 0}$, and $g$ is a filter introduced to mitigate the effect of the approximation $u \approx \tilde{v}$. It will be specified later (see Section B). Finally, $\tilde{v}_m = (v_m - \tilde{v})$ corresponds to the $m$th raw SIM data without the widefield component.

#### A. Grid-Based Initialization via Cross-correlation

Ignoring the filters $h$ and $g$, and moving to the Fourier domain, we obtain the following approximation of $\mathcal{J}$

$$\widetilde{\mathcal{J}}(\mathbf{a}, \boldsymbol{\phi}, \mathbf{k}) = \sum_{m=1}^{M} \|a_m \mathrm{e}^{\mathrm{i}\phi_m} \widetilde{V}(\cdot - \mathbf{k}) + a_m \mathrm{e}^{-\mathrm{i}\phi_m} \widetilde{V}(\cdot + \mathbf{k}) - \widetilde{V}_m\|_2^2.$$

The minimization of $\tilde{J}$ with respect to the $a_m \mathrm{e}^{\mathrm{i}\phi_m}$ amounts to minimize $M$ functions $f_m$ defined as

$$\begin{aligned} f_m(\alpha) &= \|\alpha \widetilde{V}(\cdot - \mathbf{k}) + \bar{\alpha} \widetilde{V}(\cdot + \mathbf{k}) - \widetilde{V}_m\|_2^2 \\ &= \|\alpha \widetilde{V}(\cdot - \mathbf{k})\|_2^2 + \|\bar{\alpha} \widetilde{V}(\cdot + \mathbf{k})\|_2^2 + \|\widetilde{V}_m\|_2^2 - 2\Re\left(\left\langle \alpha \widetilde{V}(\cdot - \mathbf{k}), \widetilde{V}_m \right\rangle + \left\langle \bar{\alpha} \widetilde{V}(\cdot + \mathbf{k}), \widetilde{V}_m \right\rangle\right) \\ &\qquad \cdots + 2\Re\left(\left\langle \alpha \widetilde{V}(\cdot - \mathbf{k}), \bar{\alpha} \widetilde{V}(\cdot + \mathbf{k}) \right\rangle\right). \end{aligned} \quad \textbf{(11)}$$

Because $\widetilde{V}$ has the same support as the OTF, it is compactly supported. Hence, the amplitude of the inner product $\langle \alpha \widetilde{V}(\cdot - \mathbf{k}), \bar{\alpha} \widetilde{V}(\cdot + \mathbf{k}) \rangle$ globally decreases when $\|\mathbf{k}\|$ increases. It reaches zero when $\mathbf{k}$ is outside of the OTF support. As the modulation wave vector is expected to lie close to the OTF cutoff frequency (to get the super-resolution effect), we simplify the calculations using the approximation that $\langle \alpha \widetilde{V}(\cdot - \mathbf{k}), \bar{\alpha} \widetilde{V}(\cdot + \mathbf{k}) \rangle = 0$. Then, it holds that

$$f_m(\alpha) = 2|\alpha|^2 \|\widetilde{V}\|_2^2 + \|\widetilde{V}_m\|_2^2 - 2\Re\left(\bar{\alpha}\left(\left\langle \widetilde{V}(\cdot - \mathbf{k}), \widetilde{V}_m \right\rangle + \overline{\left\langle \widetilde{V}(\cdot + \mathbf{k}), \widetilde{V}_m \right\rangle}\right)\right) \quad \textbf{(12)}$$

with $\|\widetilde{V}(\cdot \pm \mathbf{k})\|_2 = \|\widetilde{V}\|_2$ and $\Re(ab) = \Re(\bar{a}\bar{b})$ for $a, b \in \mathbb{C}$. Differentiating (12) with respect to $\alpha$, we get that

$$f'_m(\alpha) = 4\alpha \|\widetilde{V}\|_2^2 - 2\left(\left\langle \widetilde{V}(\cdot - \mathbf{k}), \widetilde{V}_m \right\rangle + \overline{\left\langle \widetilde{V}(\cdot + \mathbf{k}), \widetilde{V}_m \right\rangle}\right), \quad \textbf{(13)}$$

and thus the desired solution, denoted $\alpha_m(\mathbf{k})$ as it depends on $\mathbf{k}$ and $m$, is given by

$$f'_m(\alpha_m(\mathbf{k})) = 0 \;\Rightarrow\; \alpha_m(\mathbf{k}) = \frac{\left\langle \widetilde{V}(\cdot - \mathbf{k}), \widetilde{V}_m \right\rangle + \overline{\left\langle \widetilde{V}(\cdot + \mathbf{k}), \widetilde{V}_m \right\rangle}}{2\|\widetilde{V}\|_2^2}. \quad \textbf{(14)}$$

Injecting now this solution in (12), summing over $m$, and ignoring the constant term $\|\widetilde{V}_m\|_2^2$, we obtain an expression of $\tilde{\mathcal{J}}$ that depends only on $\mathbf{k}$, as in

$$\begin{aligned}\tilde{J}(\mathbf{k}) &= \sum_{m=1}^{M} f_m(\alpha_m(\mathbf{k})) = \sum_{m=1}^{M} 2|\alpha_m(\mathbf{k})|^2\|\widetilde{V}\|_2^2 - 2\Re\left(\overline{\alpha_m(\mathbf{k})}\alpha_m(\mathbf{k})\right)2\|\widetilde{V}\|_2^2 = -\sum_{m=1}^{M} 2|\alpha_m(\mathbf{k})|^2\|\widetilde{V}\|_2^2 \\ &= -\sum_{m=1}^{M}\frac{\left|\left\langle \widetilde{V}(\cdot-\mathbf{k}),\widetilde{V}_m\right\rangle+\overline{\left\langle \widetilde{V}(\cdot+\mathbf{k}),\widetilde{V}_m\right\rangle}\right|^2}{2\|\widetilde{V}\|_2^2}. \end{aligned} \tag{15}$$

We evaluate the function $\tilde{J}$ in (15) on a grid $\mathcal{K} = \{\mathbf{k}_q\}_{q=1}^{Q}$ of candidate wave vectors through $M$ cross-correlations between $\tilde{V}$ and $\tilde{V}_m$. This can be performed efficiently using the fast Fourier transform (FFT) together with proper padding operations if the grid $\mathcal{K}$ is chosen finer than that of the raw SIM data (see Figure S4, Panel **B**). An initial wave vector $\mathbf{k}_{\text{init}} \in \mathcal{K}$ can then be easily extracted from the image $\{\tilde{\mathcal{J}}(\mathbf{k}_q)\}_{q=1}^{Q}$. In FlexSIM, the center of this image can be masked to avoid spurious global minimizers (due to the discrete grid) that may appear for very challenging data. Finally, given $\mathbf{k}_{\text{init}}$, we obtain initial estimates of $\mathbf{a}$ and $\boldsymbol{\phi}$ with (14) as $a_m = |\alpha_m(\mathbf{k}_{\text{init}})|$ and $\phi_m = \arg(\alpha_m(\mathbf{k}_{\text{init}}))$.

Although we have not assumed that phases are equally spaced, this assumption can be easily integrated to the proposed grid-based initialization (see Section C). This can help to improve the robustness to noise and model mismatch (e.g., the use of an approximated OTF).

### B. Local Off-the-Grid Refinement

The goal of the refinement step discussed in this section is to improve the initial estimation through a local optimization of $\mathcal{J}$ in (10). This allows us to obtain an estimate of the wave vector that is not constrained to be on the grid $\mathcal{K}$ used for the initialization step. Moreover, the proposed local refinement deals with the exact function $\mathcal{J}$ in (10). It therefore accounts for the filters $h$ and $g$ in (10) and does not rely on the assumption leading to (12). Besides, given the initial estimation of the wave vector $\mathbf{k}_{\text{init}}$, we define $g$ to be a modulated version of the PSF, as

$$g(\mathbf{x}) = 2\,h(\mathbf{x})\cos\left(\mathbf{k}_{\text{init}}^{\text{t}}\mathbf{x}\right). \tag{16}$$

The rationale behind this choice is illustrated in Figure S3. It is somehow related to the idea of notch filtering used in other works [22].

As opposed to the initialization step, we work here in the spatial domain. The reason is that, in (10), the dependence of $\mathcal{J}$ in $\mathbf{k}$ enjoys a closed-form expression (via the cos). This is not the case in the Fourier domain where $\mathbf{k}$ would appear as a translation parameter of $\widetilde{V}$, a quantity which is only known in practice trough noisy samples.

Using the relation $\cos(a+b) = (\cos(a)\cos(b) - \sin(a)\sin(b))$, we rewrite $\mathcal{J}$ as

$$\mathcal{J}(\mathbf{c},\mathbf{s},\mathbf{k}) = \sum_{m=1}^{M}\|h*(c_m\tilde{v}_c - s_m\tilde{v}_s)(\mathbf{k},\cdot) - g*\tilde{v}_m\|_2^2, \tag{17}$$

where $c_m = a_m\cos(\phi_m)$, $s_m = a_m\sin(\phi_m)$, and

$$\tilde{v}_c(\mathbf{k},\mathbf{x}) = \cos(\mathbf{k}^{\text{t}}\mathbf{x})\tilde{v}(\mathbf{x}), \quad \tilde{v}_s(\mathbf{k},\mathbf{x}) = \sin(\mathbf{k}^{\text{t}}\mathbf{x})\tilde{v}(\mathbf{x}). \tag{18}$$

As such, for a fixed $\mathbf{k}$, the minimization of $\mathcal{J}$ with respect to $\mathbf{c}$ and $\mathbf{s}$ amounts to the solution $M$ systems of equations in two variables of the form $\mathbf{A}\mathbf{z}_m = \mathbf{b}_m$, where $\mathbf{z}_m = (c_m, s_m)^{\text{t}}$, $\mathbf{b}_m = (\langle h*\tilde{v}_c(\mathbf{k},\cdot), g*\tilde{v}_m\rangle, -\langle h*\tilde{v}_s(\mathbf{k},\cdot), g*\tilde{v}_m\rangle)^{\text{t}}$, and

$$\mathbf{A} = \begin{pmatrix} \|h*\tilde{v}_c(\mathbf{k},\cdot)\|_2^2 & -\langle h*\tilde{v}_c(\mathbf{k},\cdot), h*\tilde{v}_s(\mathbf{k},\cdot)\rangle \\ -\langle h*\tilde{v}_s(\mathbf{k},\cdot), h*\tilde{v}_c(\mathbf{k},\cdot)\rangle & \|h*\tilde{v}_s(\mathbf{k},\cdot)\|_2^2 \end{pmatrix}. \tag{19}$$

In contrast, the minimization of $\mathcal{J}$ in (10) with respect to $\boldsymbol{\phi}$ would involve the resolution of $M$ difficult nonlinear systems of equations. This highlights the interest of (17).

As it turns out, for fixed $\mathbf{c}$ and $\mathbf{s}$, we have access to a closed-form expression of the gradient of $\mathcal{J}$ in (17) with respect to $\mathbf{k}$. It reads

$$\boldsymbol{\nabla}_{\mathbf{k}}\mathcal{J}(\mathbf{c},\mathbf{s},\mathbf{k}) = \sum_{m=1}^{M}\begin{pmatrix} 2\langle h*\partial_{k_1}(c_m\tilde{v}_c - s_m\tilde{v}_s)(\mathbf{k},\cdot), r_m\rangle \\ 2\langle h*\partial_{k_2}(c_m\tilde{v}_c - s_m\tilde{v}_s)(\mathbf{k},\cdot), r_m\rangle \end{pmatrix} \tag{20}$$

where $r_m = (h*(c_m\tilde{v}_c - s_m\tilde{v}_s)(\mathbf{k},\cdot) - g*\tilde{v}_m)$ and

$$\partial_{k_i}(c_m\tilde{v}_c - s_m\tilde{v}_s)(\mathbf{k},\mathbf{x}) = -x_i(c_m\tilde{v}_s + s_m\tilde{v}_c)(\mathbf{k},\mathbf{x}), \tag{21}$$

for all $i = 1, 2$. Hence, we can deploy gradient descent steps to minimize $\mathcal{J}$ with respect to $\mathbf{k}$. More precisely, we perform $\mathbf{k}_{i+1} = (\mathbf{k}_i - \gamma_i \boldsymbol{\nabla}_{\mathbf{k}} \mathcal{J}(\mathbf{c}, \mathbf{s}, \mathbf{k}_i))$ for $\gamma_i > 0$ obtained through backtracking line search.

Here again, the constraint of equally spaced phases can be easily introduced as detailed in Section C).

#### C. Extension to Equally Spaced Phases

In Sections A and B, we presented our proposed pattern-estimation method in its most general form, without any assumption on the distribution of pattern phases. However, in practice, it is very common to consider the equally spaced phases $\phi_m = \phi_{\text{off}} + 2\pi(m-1)/M$. As such, the estimation of the $M$ phases $\phi_m$ is reduced to the estimation of just one phase offset $\phi_{\text{off}}$. Our pattern-estimation method can be easily modified so as to account for this.

- *Grid-Based Initialization via Cross-correlation.* Here, we need to minimize $\tilde{J}$ with respect to the single complex number $a\mathrm{e}^{\mathrm{i}\phi_{\text{off}}}$. Similarly to (12), we get that this amounts to the minimization of a single function $f$ (not depending on $m$ anymore) defined by

$$f(\alpha) = 2|\alpha|^2 \|\widetilde{V}\|_2^2 + \sum_{m=1}^{M} \|\widetilde{V}_m\|_2^2 - 2\Re\left(\bar{\alpha}\mathrm{e}^{-\mathrm{i}\frac{2\pi(m-1)}{M}}\left(\left\langle \widetilde{V}(\cdot - \mathbf{k}), \widetilde{V}_m \right\rangle + \overline{\left\langle \widetilde{V}(\cdot + \mathbf{k}), \widetilde{V}_m \right\rangle}\right)\right). \quad \textbf{(22)}$$

This function is minimized for

$$\alpha(\mathbf{k}) = \sum_{m=1}^{M} \mathrm{e}^{-\mathrm{i}\frac{2\pi(m-1)}{M}} \frac{\left\langle \widetilde{V}(\cdot - \mathbf{k}), \widetilde{V}_m \right\rangle + \overline{\left\langle \widetilde{V}(\cdot + \mathbf{k}), \widetilde{V}_m \right\rangle}}{2\|\widetilde{V}\|_2^2}, \quad \textbf{(23)}$$

which is the counterpart of (14) when equally spaced phases are assumed. Injecting this solution in $f$ leads to

$$\tilde{J}(\mathbf{k}) = -2|\alpha(\mathbf{k})|^2 \|\widetilde{V}\|_2^2 = -\left| \sum_{m=1}^{M} \mathrm{e}^{-\mathrm{i}\frac{2\pi(m-1)}{M}} \frac{\left\langle \widetilde{V}(\cdot - \mathbf{k}), \widetilde{V}_m \right\rangle + \overline{\left\langle \widetilde{V}(\cdot + \mathbf{k}), \widetilde{V}_m \right\rangle}}{2\|\widetilde{V}\|_2^2} \right|^2 \quad \textbf{(24)}$$

which can be computed efficiently for all $\mathbf{k} \in \mathcal{K}$ through the same $M$ cross-correlation maps as for the case of uneven phase shifts.

- *Local Off-the-Grid Refinement.* Under the assumption of equally spaced phase shifts (and $a_m = a$ for all $m$), (17) becomes

$$\mathcal{J}(c, s, \mathbf{k}) = \sum_{m=1}^{M} \|h * (c\, \tilde{v}_{c,m} - s\, \tilde{v}_{s,m})(\mathbf{k}, \cdot) - g * \tilde{v}_m\|_2^2, \quad \textbf{(25)}$$

where $c = a\cos(\phi_{\text{off}})$, $s = a\sin(\phi_{\text{off}})$, and

$$\tilde{v}_{c,m}(\mathbf{k}, \mathbf{x}) = \cos\left(\mathbf{k}^{\mathrm{t}}\mathbf{x} + 2\pi(m-1)/M\right) \tilde{v}(\mathbf{x}), \qquad \tilde{v}_{s,m}(\mathbf{k}, \mathbf{x}) = \sin\left(\mathbf{k}^{\mathrm{t}}\mathbf{x} + 2\pi(m-1)/M\right) \tilde{v}(\mathbf{x}). \quad \textbf{(26)}$$

Then, for a fixed $\mathbf{k}$, the minimization of $\mathcal{J}$ with respect to $c$ and $s$ is now restricted to the resolution of the single $2 \times 2$ system of linear equations

$$\sum_{m=1}^{M} \begin{pmatrix} \|h * \tilde{v}_{c,m}(\mathbf{k}, \cdot)\|_2^2 & -\langle h * \tilde{v}_{c,m}(\mathbf{k}, \cdot), h * \tilde{v}_{s,m}(\mathbf{k}, \cdot)\rangle \\ -\langle h * \tilde{v}_{s,m}(\mathbf{k}, \cdot), h * \tilde{v}_{c,m}(\mathbf{k}, \cdot)\rangle & \|h * \tilde{v}_{s,m}(\mathbf{k}, \cdot)\|_2^2 \end{pmatrix} \begin{pmatrix} c \\ s \end{pmatrix} = \sum_{m=1}^{M} \begin{pmatrix} \langle h * \tilde{v}_{c,m}(\mathbf{k}, \cdot), g * \tilde{v}_m\rangle \\ -\langle h * \tilde{v}_{s,m}(\mathbf{k}, \cdot), g * \tilde{v}_m\rangle \end{pmatrix}. \quad \textbf{(27)}$$

Concerning the update of $\mathbf{k}$ through gradient-descent steps, the consideration of equally spaced phase shifts only changes the gradient. It becomes

$$\boldsymbol{\nabla}_{\mathbf{k}} \mathcal{J}(c, s, \mathbf{k}) = \sum_{m=1}^{M} \begin{pmatrix} 2\,\langle h * \partial_{k_1}(c\, \tilde{v}_{c,m} - s\, \tilde{v}_{s,m})(\mathbf{k}, \cdot), r_m\rangle \\ 2\,\langle h * \partial_{k_2}(c\, \tilde{v}_{c,m} - s\, \tilde{v}_{s,m})(\mathbf{k}, \cdot), r_m\rangle \end{pmatrix}, \quad \textbf{(28)}$$

where $r_m = (h * (c\, \tilde{v}_{c,m} - s\, \tilde{v}_{s,m})(\mathbf{k}, \cdot) - g * \tilde{v}_m)$ and $\partial_{k_i}(c\, \tilde{v}_{c,m} - s\, \tilde{v}_{s,m})(\mathbf{k}, \mathbf{x}) = -x_i(c\, \tilde{v}_{s,m} + s\, \tilde{v}_{c,m})(\mathbf{k}, \mathbf{x})$.

### Supplementary Note 2. Experiments on Simulated Data

In this note we provide some quantitative comparisons of FlexSIM with existing methods to complement the extensive comparisons reported on the real data of Table S2.

#### A. Simulation Settings

We consider the test phantom presented in Figure S1 which contains a variety of structures (lines, dots, flat regions, start-like object). Illumination patterns are generated according to the two-beam model

$$w(\mathbf{x}) = 1 + a\cos(\mathbf{k}^t\mathbf{x} + \phi), \quad \textbf{(29)}$$

where the modulation contrast is fixed to $a = 0.5$ and, for a given pattern orientation, three phases shifts $\phi$ spaced by $2\pi/3$ are considered. For each pattern orientation $\theta \in (0, 2\pi]$, the corresponding wavevector is computed as $\mathbf{k} = k\sin(\beta)(\cos(\theta), \sin(\theta))^t$ where $k = 2\pi n_{\text{sam}}/\lambda_{\text{exc}}$, with $\lambda_{\text{exc}} = 520$ nm the excitation wavelength and $n_{\text{sam}} = 1.333$ the refractive index of the sample, and where $\beta = 80°$ denotes the angle between the optical axis and interfering beams. The pixel size (of acquisition grid) is set to 84 nm. With these parameters, the sampling frequency is slightly above the Nyquist rate and patterns frequencies are close to the OTF cutoff so as to maximize the super-resolution effect. Finally, a background of 10% of the maximum intensity is added to the SIM data which are then corrupted with a Poisson noise corresponding to a maximum expected number of photons of 100.

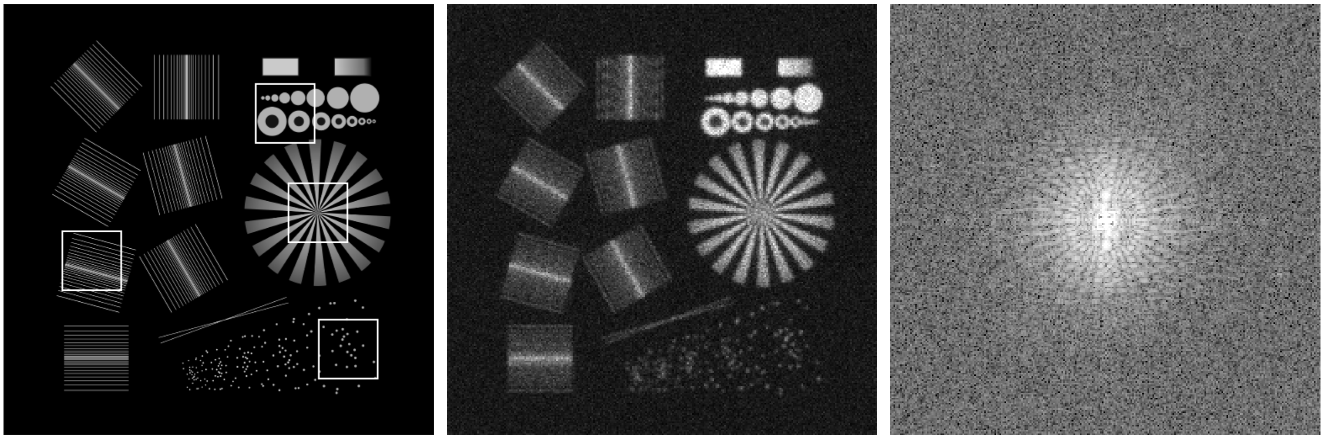

**Fig. S1. Simulated dataset.** From left to right: synthetic phantom from [59], example of SIM data and its Fourier transform.

#### B. Reconstruction Results

From these data, we reconstructed a super-resolved image using FairSIM [11, 18], OpenSIM [19], Direct-SIM [25], PCA-SIM [63], HiFi-SIM [22], JSFR-SIM [24] and the proposed FlexSIM. These correspond to all the methods reported in Table S2 expect ML-SIM [23] for which we couldn't manage to get a reconstruction for this simulated dataset. Yet, note that the superiority of FlexSIM on ML-SIM is clear from Figures 4.A and S11. We tuned the parameters of each method so as to maximize its performance. They are given respectively by

- FairSIM: Wiener parameter = 0.45 / APObend = 1.9,
- OpenSIM: No parameters available to be tuned, they are computed automatically,
- Direct-SIM: $\alpha$=0.5, $\beta$=0.1, ApoFHWM=0.3,
- PCASIM: No parameter for the pattern estimation (PCA related) and reconstruction is done with HiFi-SIM (see below),
- HiFi-SIM: ApoFHWM=0.31, $\beta$=2.3, $w_1$=1.3,
- JSFR-SIM: Wiener parameter = 0.57,
- FlexSIM: $\mu = 5 \cdot 10^{-5}$ (regularization parameter Tikhonov order 1).

We report in Table S1 peak signal-to-noise ratio (PSNR) and structured similarity index (SSIM) metrics. Zooms of obtained reconstructions are presented in Figure S2. We observe that FlexSIM leads to the best reconstruction. For this "standard" dataset (no illumination distortion), we mainly attribute this improvement to the fact that FlexSIM uses a gradient-based regularizer (order-1 Tikhonov) while other methods relies on Wiener based regularization.

**Table S1. PSNR and SSIM metrics for the simulated dataset of Figure S1.** The best score is highlighted in bold while the second best is underlined. Zooms of the corresponding reconstructions are shown in Figure S2.

| | FairSIM | OpenSIM | Direct-SIM | PCA-SIM | HiFi-SIM | JSFR-SIM | FlexSIM |
|---|---|---|---|---|---|---|---|
| PSNR (dB) | 21.15 | 20.99 | 20.9 | 21.17 | <u>21.32</u> | 21 | **22.11** |
| SSIM | 0.48 | 0.5 | 0.52 | <u>0.66</u> | 0.65 | 0.51 | **0.7** |

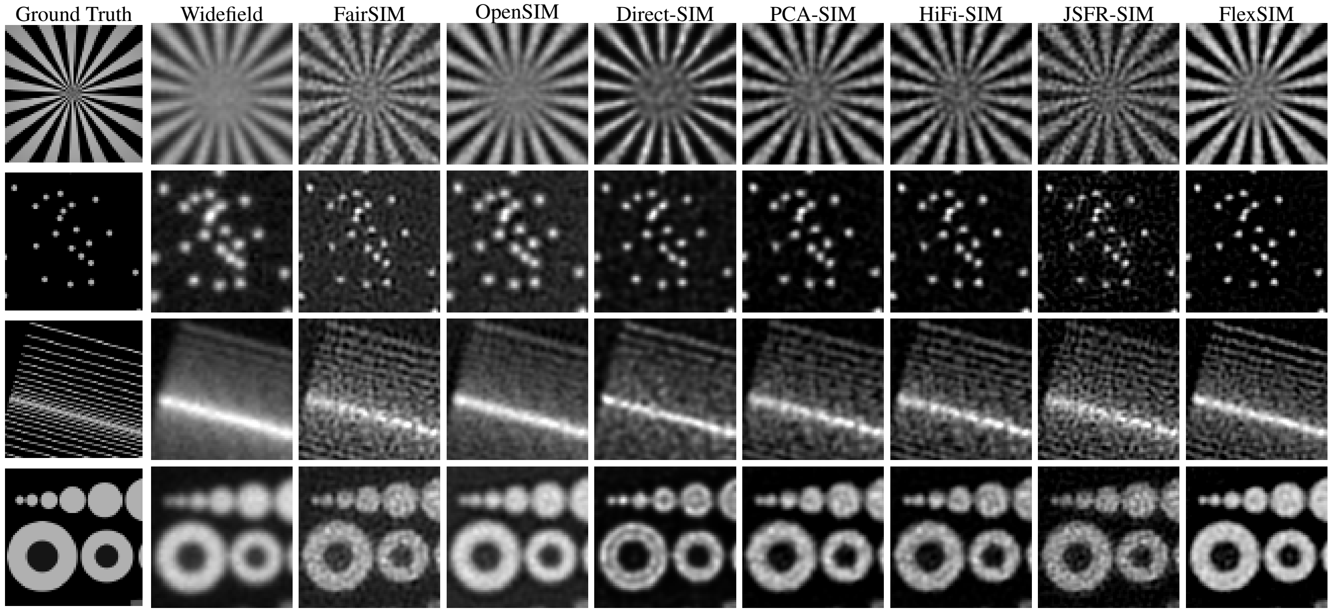


**Fig. S2. Comparisons of FlexSIM with SOTA methods on simulated data.** Zooms of the reconstructions obtained for the simulated dataset of Figure S1. Corresponding PSNR and SSIM metrics are provided in Table S1.

**Table S2. Datasets available on the web.** For each dataset we provide, the reference and reconstruction method along which it has been released, the imaged biological structures, the number of orientations (Nb orr) and phases (Nb ph), the emission wavelength ($\lambda_{\mathrm{em}}$), the numerical aperture (NA), the size of raw data, and the figure where a comparison with FlexSIM is reported.

| Reference | Biological structure | | Nb orr | Nb ph | $\lambda_{\mathrm{em}}$ | NA | Size (Raw) | Figure |
|---|---|---|---|---|---|---|---|---|
| FairSIM [11, 18] | Tubulin | | 3 | 3 | 525 | 1.49 | $256 \times 256$ | Fig. S10.A |
| | Tetraspeck beads | | 4 | 3 | 680 | 1.2 | $512 \times 512$ | Fig. S10.B |
| OpenSIM [19] | Microtubules | | 3 | 3 | 515 | 1.49 | $512 \times 512$ | Fig. S10.C |
| ML-SIM [23] | Microtubules | | 3 | 3 | 530 | 1.2 | $512 \times 512$ | Fig. 4.A |
| | Endoplasmic reticulum | | 3 | 3 | 507 | 1.2 | $512 \times 512$ | Fig. S11.A |
| | Membrane | | 3 | 3 | 530 | 1.2 | $512 \times 512$ | Fig. S11.B |
| | Beads | | 3 | 3 | 567 | 1.2 | $512 \times 512$ | Fig. S11.C |
| HiFi-SIM [22] | Microtubules | | 3 | 3 | 525 | 1.42 | $512 \times 512$ | Fig. 4.B |
| JSFR-SIM [24] | Microtubules | | 3 | 3 | 520 | 1.49 | $512 \times 512$ | Fig. 4.C |
| Direct-SIM [25] | Microtubules | | 3 | 3 | 527 | 1.42 | $512 \times 512$ | Fig. 4.D |
| | Argolight slide | | 3 | 3 | 527 | 1.42 | $512 \times 512$ | Fig. S12.A |
| | Microtubules | | 3 | 3 | 525 | 1.49 | $512 \times 512$ | Fig. S12.B |
| | Mitochondria | | 3 | 3 | 525 | 1.49 | $512 \times 512$ | Fig. S12.C |
| PCA-SIM [63] | Actin | | 3 | 3 | 607 | 1.42 | $1024 \times 1024$ | Fig. 4.E |
| | Actin | Cell 1 | 3 | 3 | 607 | 1.4 | $1024 \times 1024$ | Fig. S13.A |
| | Mitochondria | Cell 1 | 3 | 3 | 525 | 1.4 | $1024 \times 1024$ | Fig. S13.A |
| | Nucleus | Cell 1 | 3 | 3 | 460 | 1.4 | $1024 \times 1024$ | Fig. S13.A |
| | Actin | Cell 2 | 3 | 3 | 607 | 1.4 | $1024 \times 1024$ | Fig. S13.B |
| | Mitochondria | Cell 2 | 3 | 3 | 525 | 1.4 | $1024 \times 1024$ | Fig. S13.B |
| | Nucleus | Cell 2 | 3 | 3 | 460 | 1.4 | $1024 \times 1024$ | Fig. S13.B |

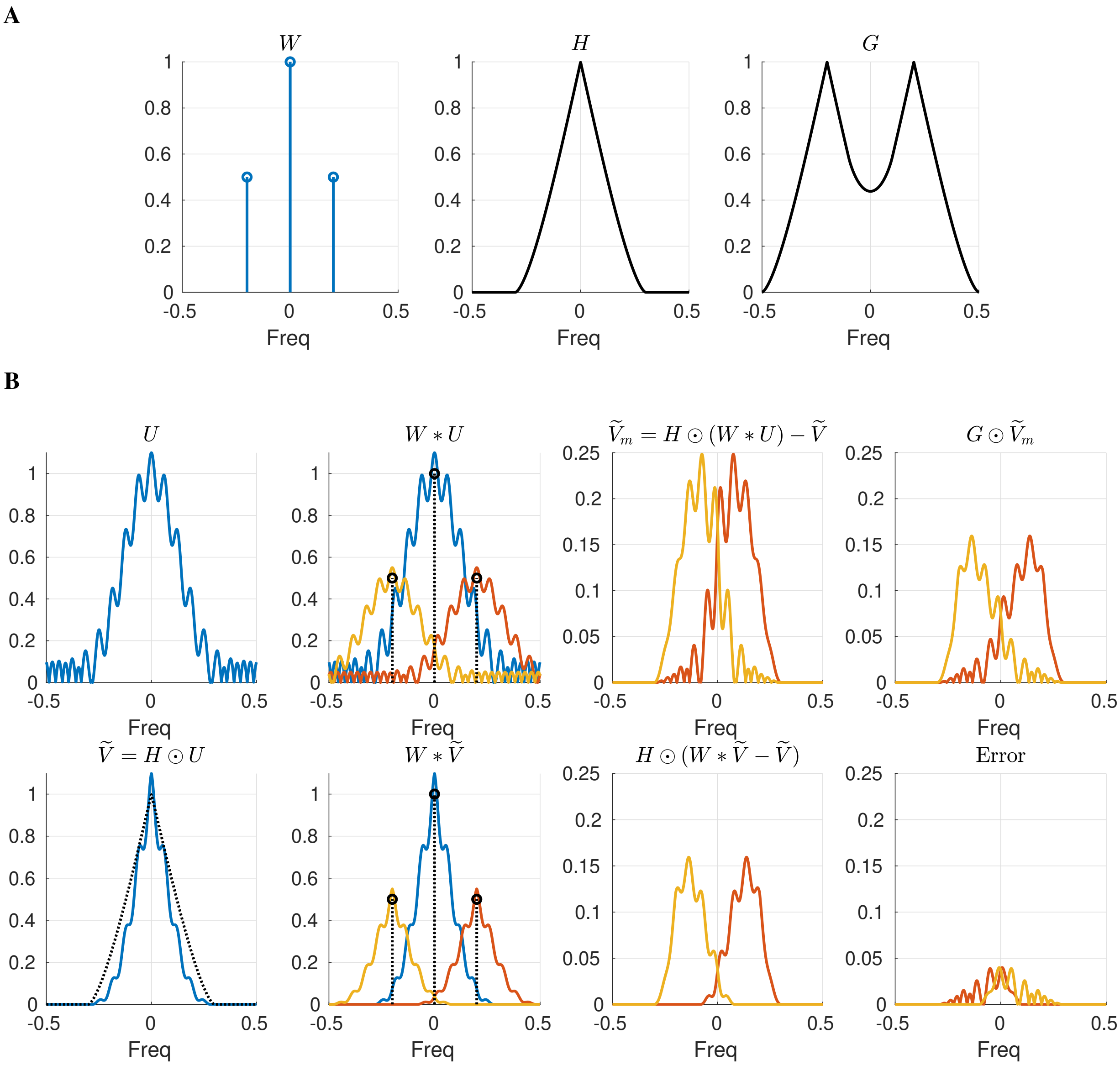


**Fig. S3. Effect of the filter $g$ in (4). A** 1D Fourier transforms of a sinusoidal pattern ($W$), an OTF ($H$), and the associated filter defined in (16) ($G$). **B**, Effect of the approximation $u \approx \tilde{v}$ we made to define $\mathcal{J}$ in (4). The different Fourier components of the modulated signals are distinguished by their color. A comparison of the two graphs in the third column reveals a mismatch between the pre-processed data $\tilde{v}_m$ (with Fourier transform $\widetilde{V}_m$) and the considered model in (4), i.e., $h * (\cos(\mathbf{k}^{\mathrm{t}} \cdot + \phi_m) \odot \tilde{v})$ (with Fourier transform $H \odot (W * \widetilde{V} - \widetilde{V})$). In contrast, the filtered data $g * \tilde{v}_m$ (with Fourier transform $G \odot \widetilde{V}_m$) is significantly more consistent with this model as demonstrated with the error plot in the bottom right graph..

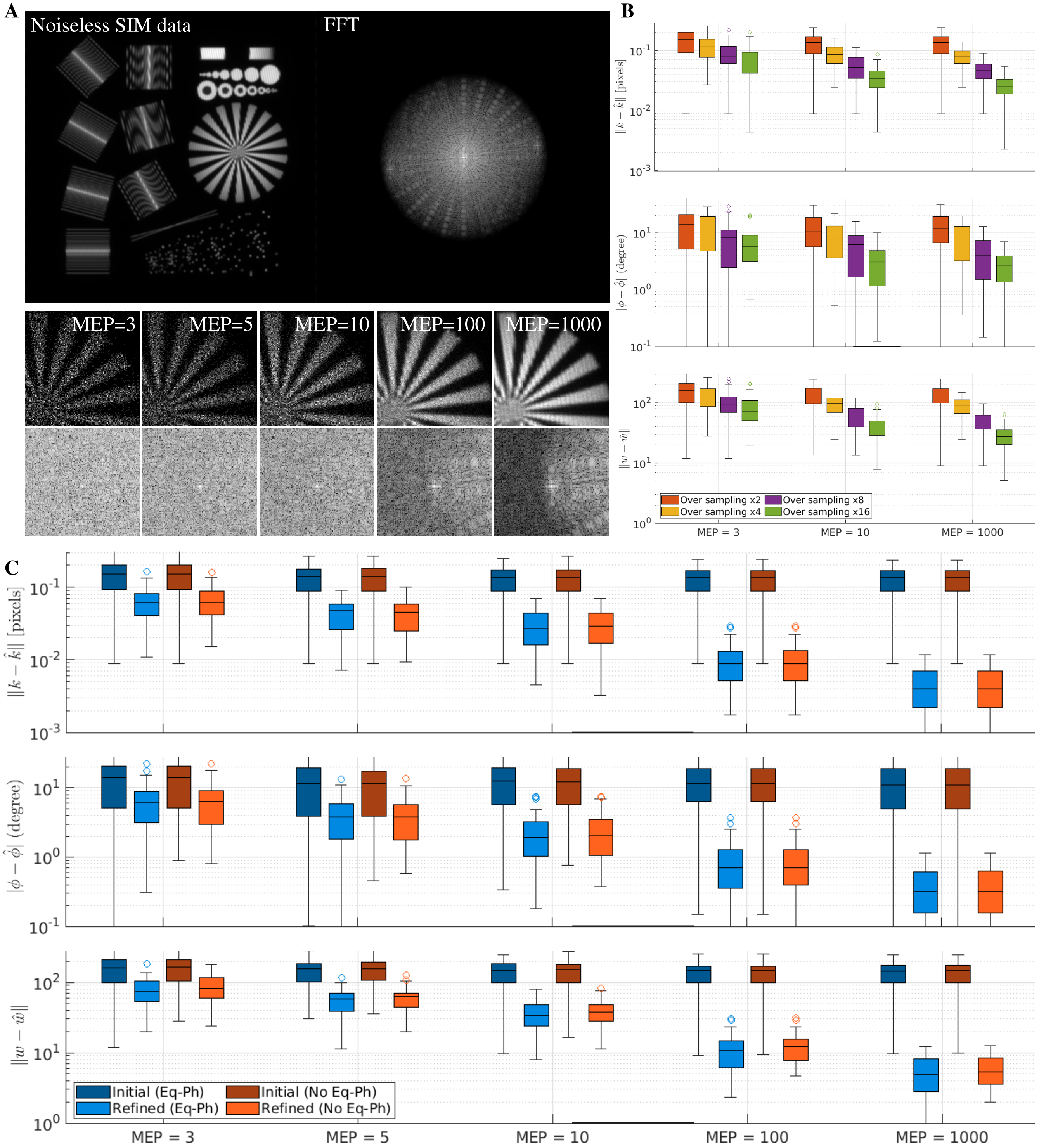


**Fig. S4. Numerical validation of FlexSIM patterns estimation.** We simulated $50$ SIM datasets with randomly generated orientations and phases using the phantom proposed in [59] (Panel **A**, top row). Simulation parameters have been set as follows: $\mathrm{NA} = 1.2$, $\lambda_{\mathrm{em}} = 530\,\mathrm{nm}$, pixel size of $64\,\mathrm{nm}$, amplitude of patterns $a = 0.9$. We then corrupted each simulated dataset with different levels of Poisson noise by varying the maximum expected number of photons (MEP). Examples (zooms) of corrupted data and their Fourier transform are depicted in the two last rows of Panel **A**. In Panel **B**, we assess the performance of the initialization step for various MEP and oversampling factors in the computation of cross-correlation maps. These results have been obtained under the assumption of equally spaced phases. In Panel **C**, we illustrate the benefit of the proposed refinement step. More precisely, we compare estimation errors for various MEP, before and after refinement, as well as with and without the assumption that the phases are equally spaced. Here the oversampling factor in the computation of cross-correlation maps (initialization step) was fixed to 2. In panels **B** and **C** the three graphs correspond to the metrics (from top to bottom): wave vector error, phase error, and the error on the patterns generated from the estimated parameters.

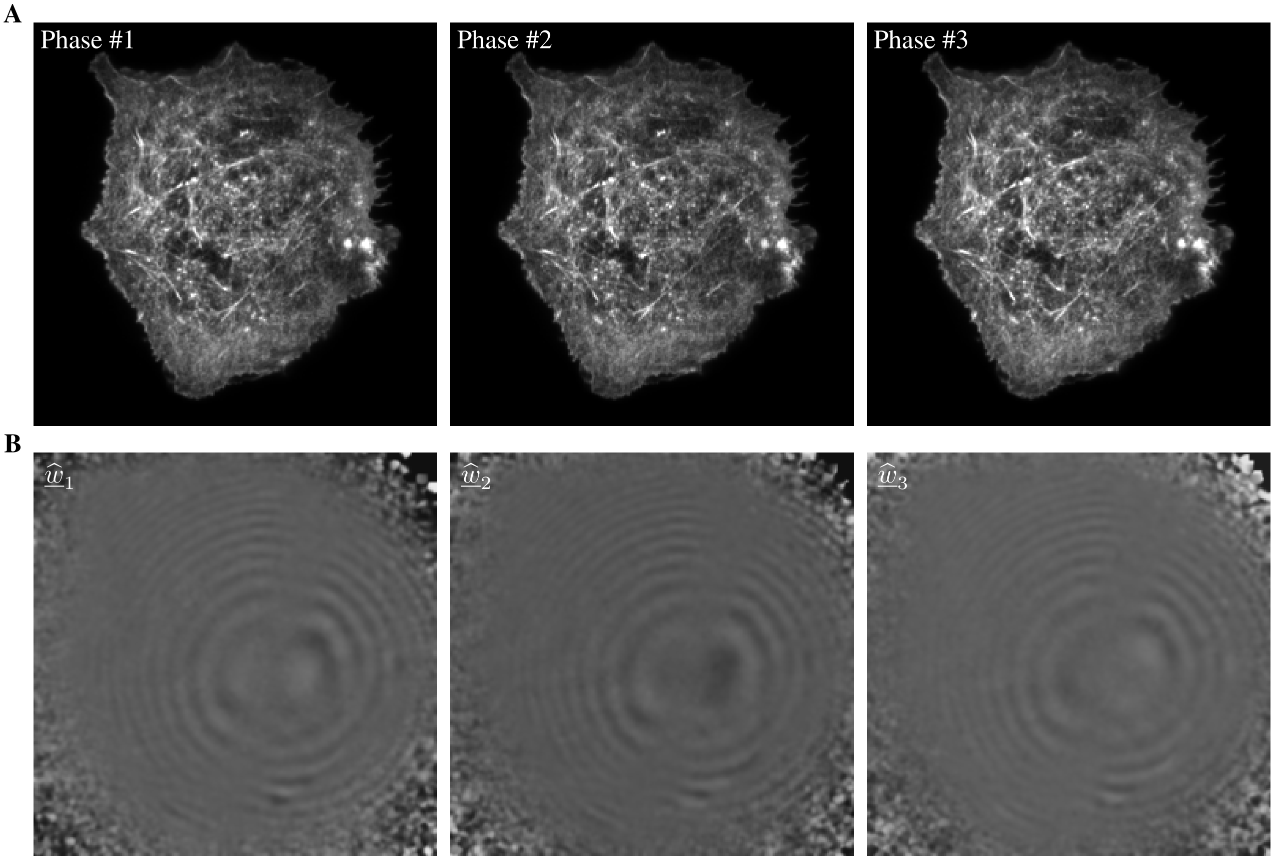


**Fig. S5. Estimation of pattern low-frequency component. A**) Raw TIRF-SIM data of COS-7 cells acquired with the Nikon N-SIM-S system in TIRF-SIM mode. All images have been acquired using the same pattern orientation but different phases. Note that, due to the TIRF mode, sinusoidal fringes are not visible. By switching rapidly from one image to another, we can observe the presence of slowly varying concentric rings. These are not clearly apparent when observing the images individually as in Panel **A**. However, as shown in Panel **B**, they are perfectly unveiled by the proposed method to estimate the low-frequency component of the pattern (see Eq. (5)).

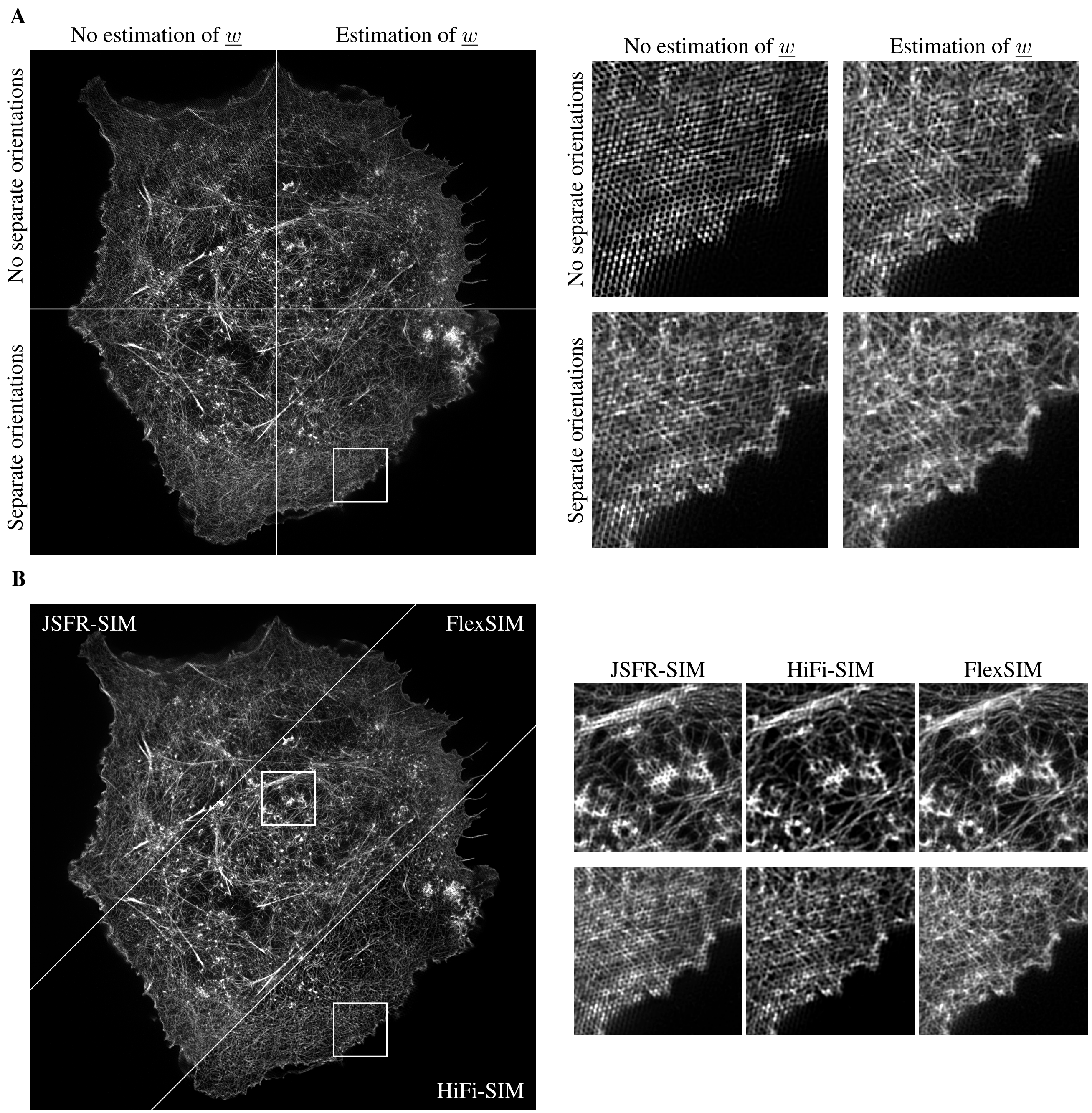


**Fig. S6. Importance of the low-frequency pattern component $\underline{w}$ and of the reconstruction of each orientation separately for challenging TIRF-SIM data.** In Panel **A**, we compare FlexSIM reconstructions obtained with and without the estimation of $\underline{w}$ as well as with and without reconstructing each orientation separately. Clearly the activation of both options is crucial to avoid reconstruction artifacts. In Panel **B** we show that the advanced features of the state-of-the-art HiFi-SIM [22] (optimisation of the reconstruction OTF) and JSFR-SIM [24] methods also fail in dealing with these challenging data. This strengthens the relevance and importance of the new features proposed in FlexSIM. HiFi-SIM reconstruction has been obtained with parameters $\mathrm{attStrength} = 0$ , $\mathrm{ApoFWHM} = 0.4$, $\beta = 0.5$, and $w_1 = 2$. JSFR-SIM reconstruction has been obtained with parameters $\mathrm{Wiener} = 0.5$, $\mathrm{Amp} = 0.98$, and $\mathrm{Sigma} = 3.5$.

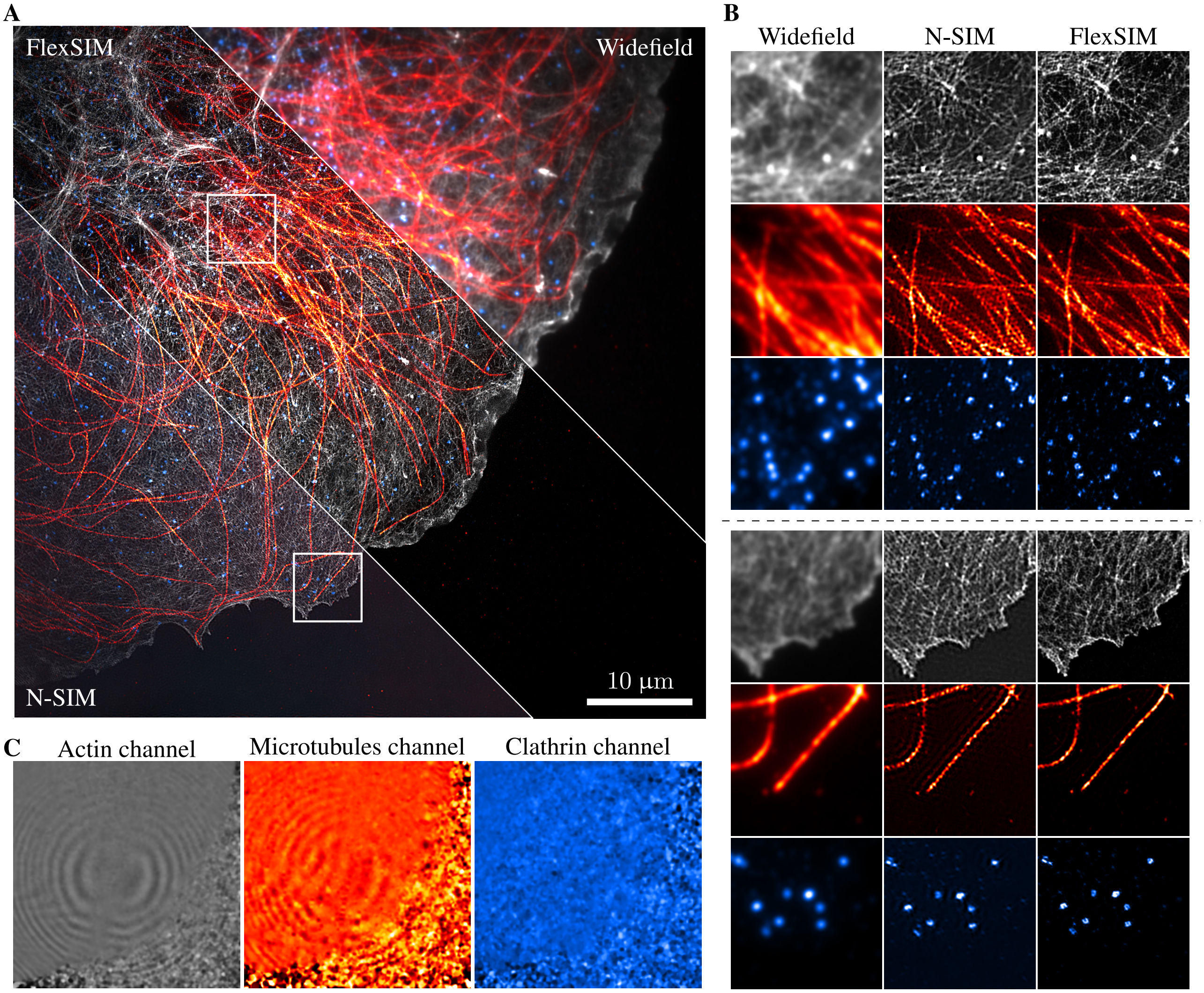


**Fig. S7. FlexSIM reconstruction of challenging TIRF-SIM data.** Actin network (gray), microtubules (red), and clathrin (blue) in COS-7 cells imaged using the Nikon N-SIM-S system in TIRF-SIM mode. Raw images are of size $(1024 \times 1024)$ and reconstructions are $(2048 \times 2048)$. **A-B)** Comparisons of reconstructions obtained with FlexSIM and the N-SIM software. As for the data reported in Figure 2, the actin N-SIM reconstruction presents grid artifacts within some regions of the sample (see second zoom in Panel **B**). In contrast, no artifacts are visible on FlexSIM reconstructions. **C)** For this sample, low frequency patterns components estimated by FlexSIM are non-uniform only for the gray and red channels. These also correspond to the channels where the N-SIM reconstruction exhibits grid artifacts.

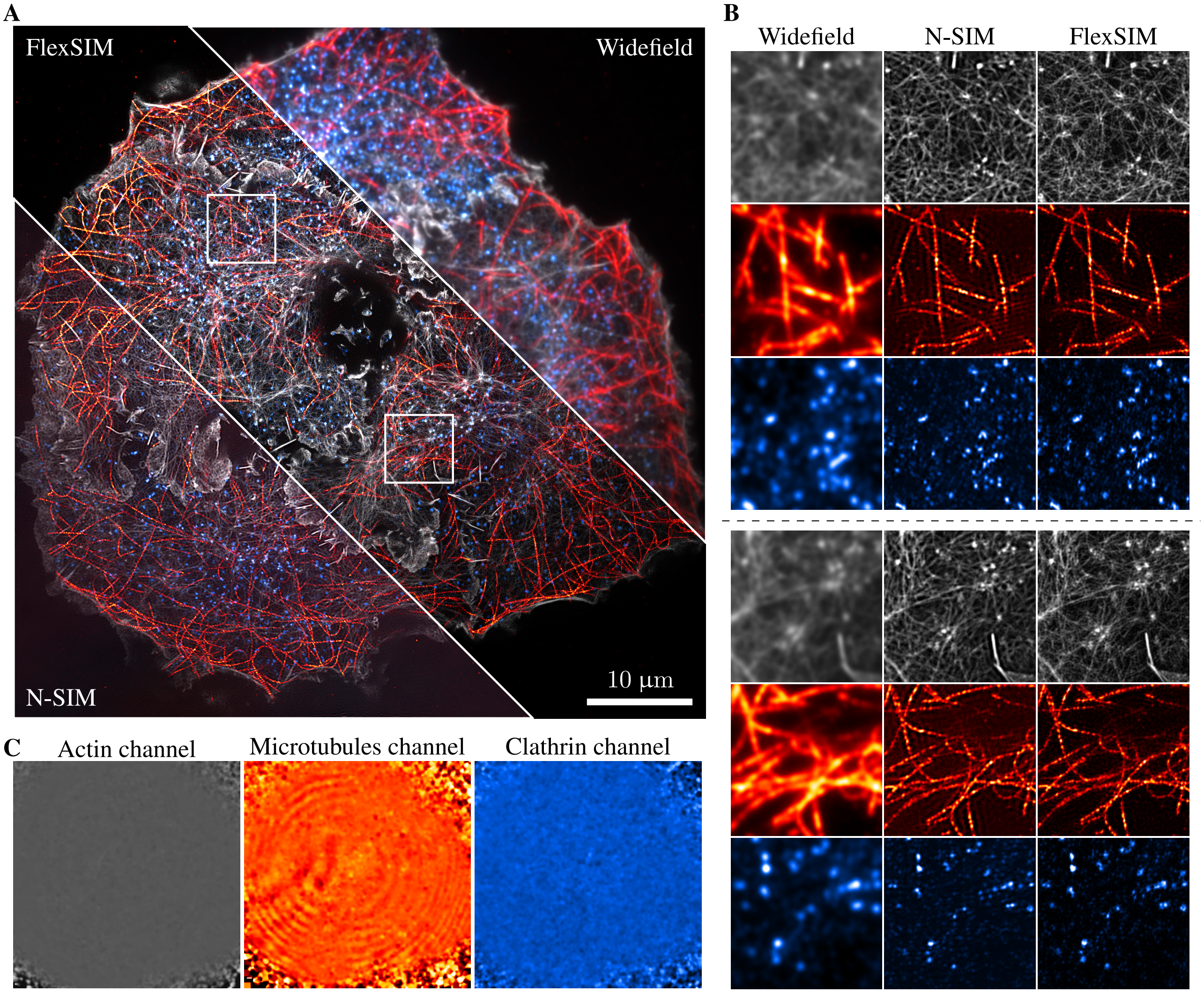


**Fig. S8. FlexSIM reconstruction of challenging TIRF-SIM data.** Actin network (gray), microtubules (red), and clathrin (blue) in COS-7 cells imaged using the Nikon N-SIM-S system in TIRF-SIM mode. Raw images are of size $(1024 \times 1024)$ and reconstructions are $(2048 \times 2048)$. **A-B)** Comparisons of reconstructions obtained with FlexSIM and the N-SIM software. As opposed to the data reported on Figures 2 and S7, here the actin N-SIM reconstruction does not present any grid artifacts. Yet, one can appreciate the better contrast and dynamic of the reconstruction obtained with FlexSIM. **C)** For this sample, a non-uniform low frequency pattern component has been estimated only for the red channel. This emphasises the fact that grid artifacts observed in N-SIM reconstructions are due to such patterns distortions. We refer the reader to Figure S6 for a further evidence of this fact.

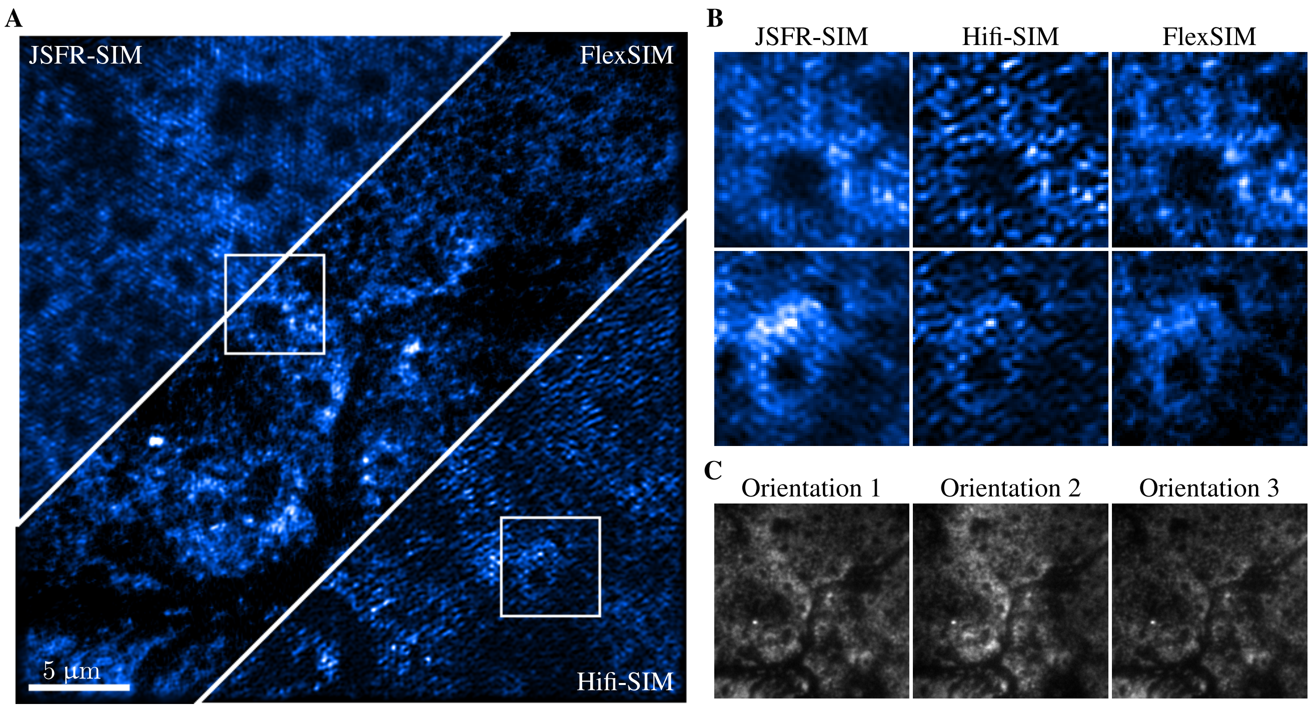


**Fig. S9. FlexSIM reconstruction of challenging TIRF-SIM data.** Imaging of paxillin in macrophage podosomes with a home-build TIRF-SIM system [35]. Raw images are of size $(256 \times 256)$ and reconstructions are $(512 \times 512)$. **A-B)** Comparisons of reconstructions obtained with FlexSIM and the HiFi-SIM and JSFR-SIM software. HiFi-SIM reconstruction has been obtained with parameters $\mathrm{attStrength} = 0$, $\mathrm{ApoFWHM} = 0.5$, $\beta = 1.2$, and $w_1 = 0.9$. JSFR-SIM reconstruction has been obtained with parameters $\mathrm{Wiener} = 0.35$, $\mathrm{Amp} = 0.98$, and $\mathrm{Sigma} = 2$. **C)** Examples of raw data for each pattern orientation which exhibit inhomogeneous illumination.

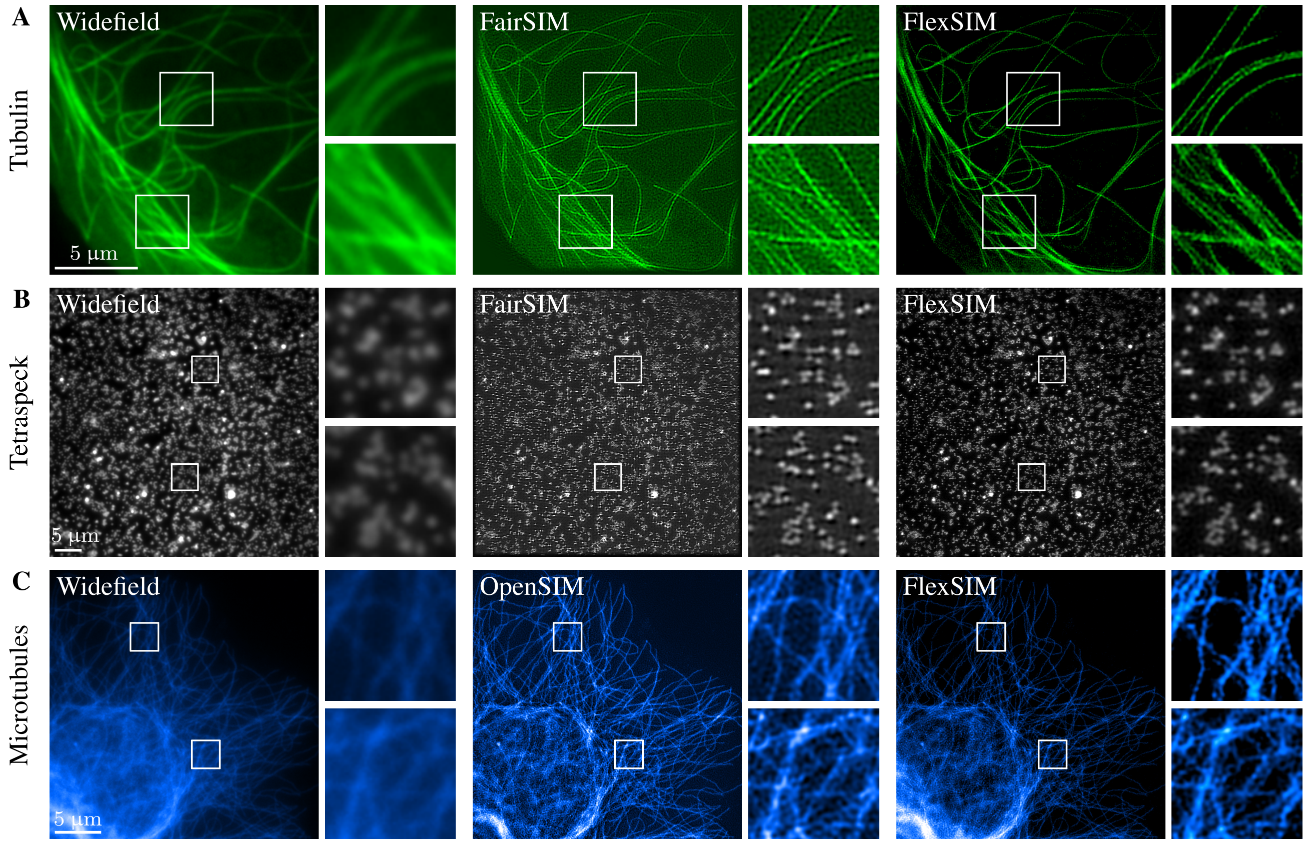


**Fig. S10. Comparisons of FlexSIM with FairSIM [18] and OpenSIM [19] on their own datasets.** Information regarding each dataset are provided in Table S2.

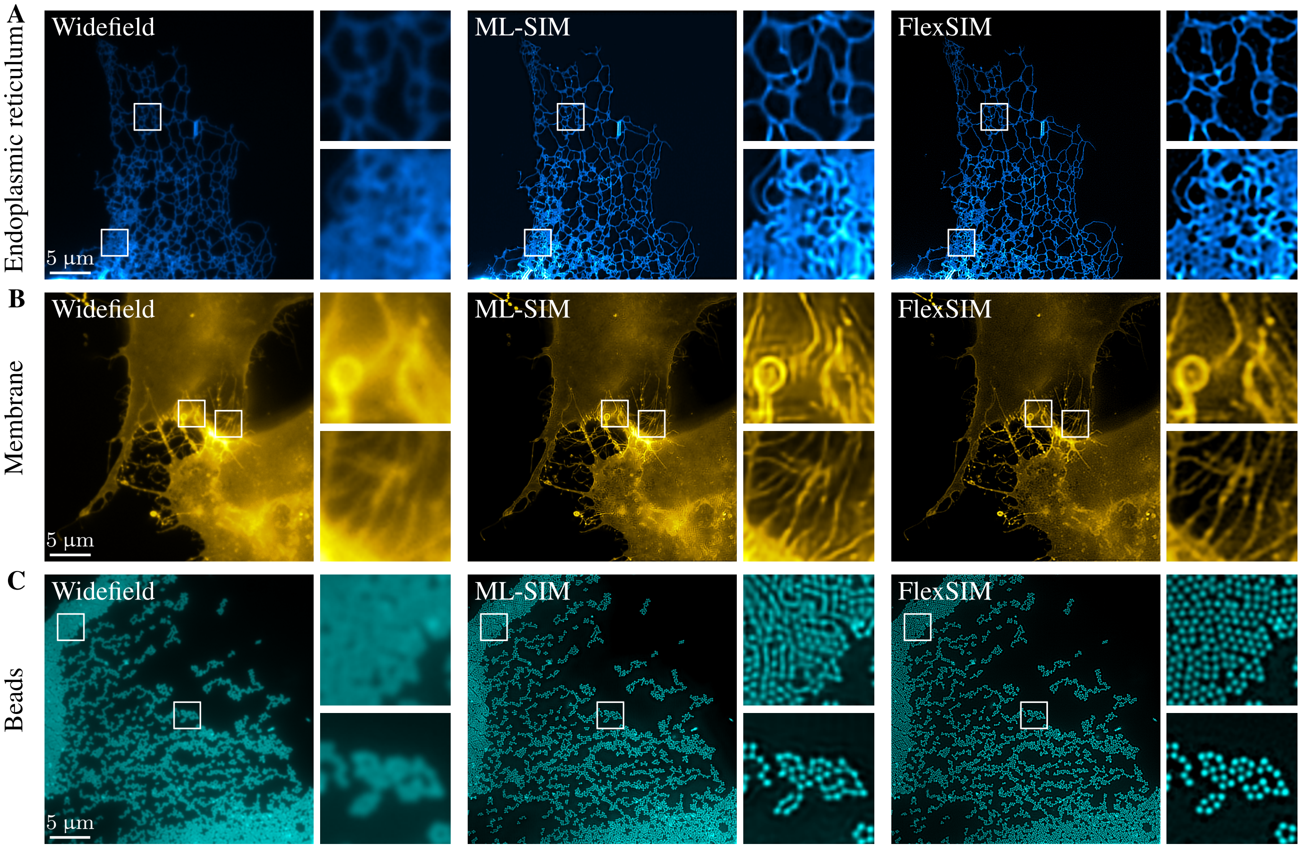


**Fig. S11. Comparisons of FlexSIM with ML-SIM [23] on its own datasets.** Information regarding each dataset are provided in Table S2. Another comparison with ML-SIM is reported on Figure 4.

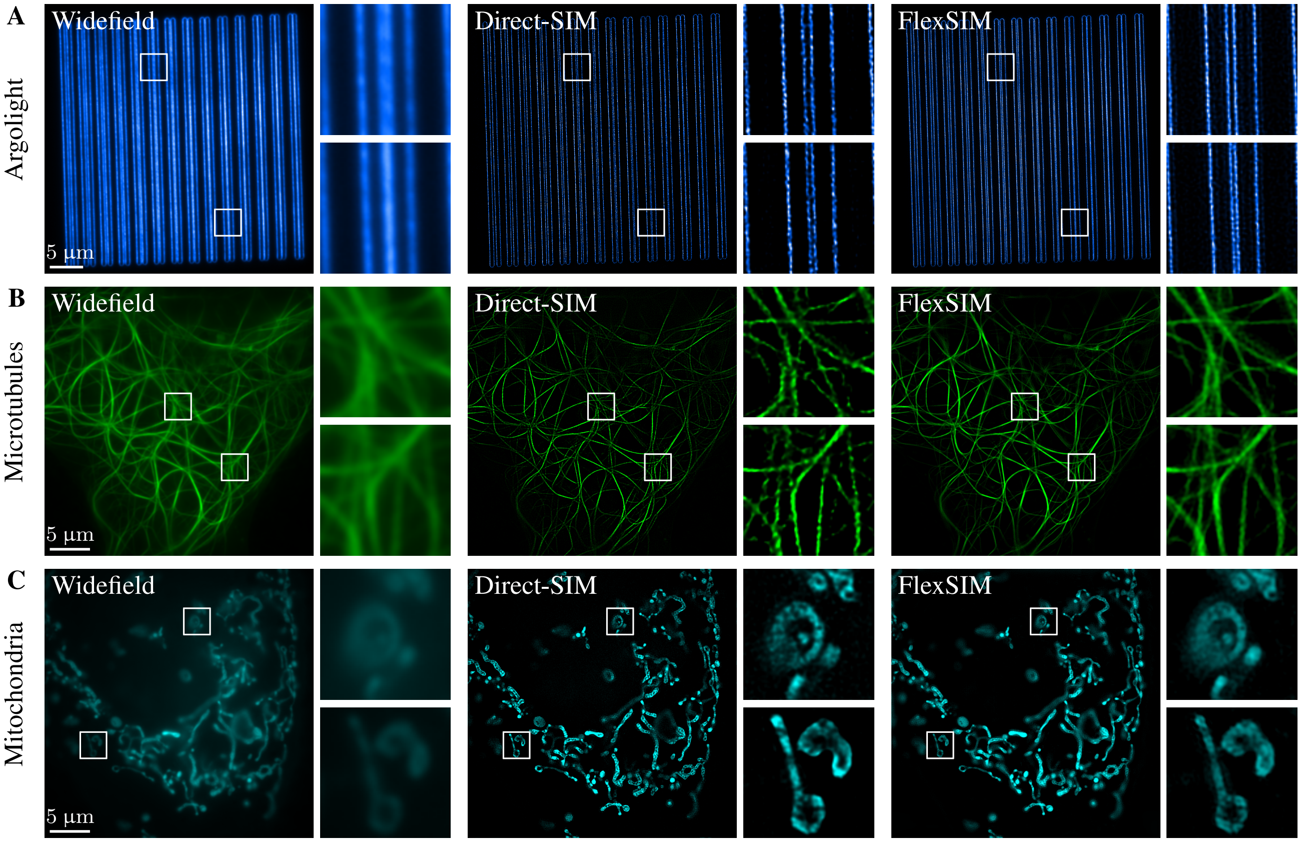


**Fig. S12. Comparisons of FlexSIM with Direct-SIM [25] on its own datasets.** Information regarding each dataset are provided in Table S2. Another comparison with Direct-SIM is reported on Figure 4.

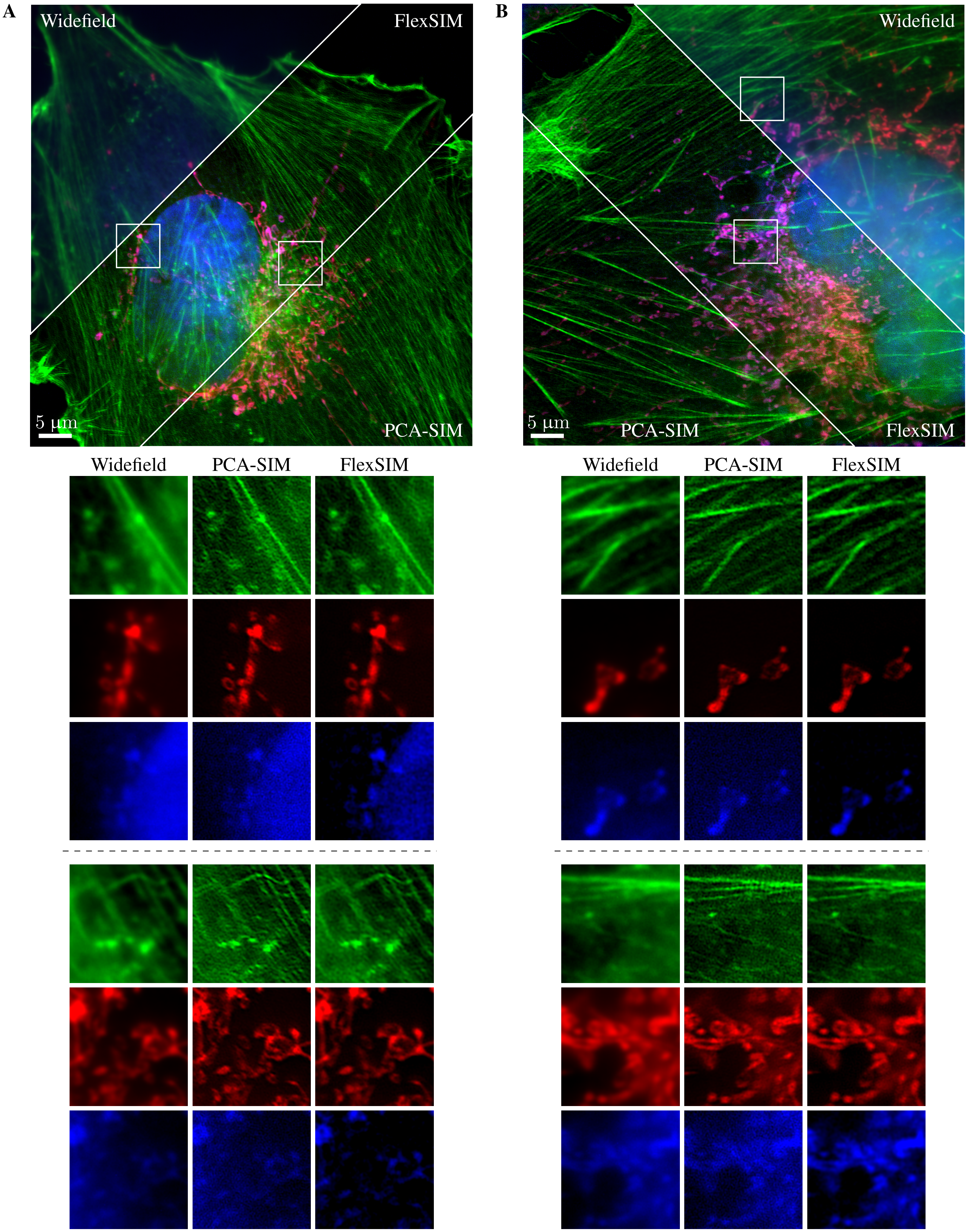


**Fig. S13. Comparisons of FlexSIM with PCA-SIM [63] on its own datasets.** Information regarding each dataset are provided in Table S2. Another comparison with PCA-SIM is reported on Figure 4. Note that PCA-SIM [63] is a method that uses principal component analysis for the estimation of patterns parameters. The reconstruction is then performed using HiFi-SIM [22].
